# Interfacial actin protrusions mechanically potentiate killing by cytotoxic T cells

**DOI:** 10.1101/443309

**Authors:** Fella Tamzalit, Mitchell S. Wang, Weiyang Jin, Vitaly Boyko, John M. Heddleston, Charles T. Black, Lance C. Kam, Morgan Huse

## Abstract

Cytotoxic T lymphocytes (CTLs) kill by forming immunological synapses with target cells and secreting toxic proteases and the pore forming protein perforin into the intercellular space. Immunological synapses are highly dynamic structures that potentiate perforin activity by applying mechanical force against the target cell. Here, we employed high-resolution imaging and microfabrication to investigate how CTLs exert synaptic forces and coordinate their mechanical output with perforin secretion. Using micropatterned stimulatory substrates that enable synapse growth in three dimensions, we found that perforin release occurs at the base of actin-rich protrusions that extend from central and intermediate locations within the synapse. These protrusions, which depended on the cytoskeletal regulator WASP and the Arp2/3 actin nucleation complex, were required for synaptic force exertion and efficient killing. They also mediated physical distortion of the target cell surface during CTL-target cell interactions. Our results reveal the mechanical basis of cellular cytotoxicity and highlight the functional importance of dynamic, three-dimensional architecture in immune cell-cell interfaces.

**One sentence summary:** Cytotoxic T lymphocytes use F-actin-rich protrusions at the immunological synapse to potentiate perforin-and granzyme-mediated target cell killing.

## INTRODUCTION

Cell-cell interactions are critical mediators of intercellular communication. They guide movement, inform cell fate, and coordinate collective behavior during development and homeostasis. They also propagate cellular dysfunction in the context of cancer and other disease states. It is thought that the structure of each cell-cell interaction is tailored precisely to its respective function. Exploring this relationship between structure and function is challenging, however, because many interactions have intricate, three-dimensional architectures that are difficult to reconstitute in controlled environments. In addition, they are often highly dynamic, necessitating the application of precise, time-resolved analytical tools to achieve mechanistic insight. Finally, experimental strategies that disrupt specific features of interfacial architecture are usually unavailable, making it difficult to ascribe function to those features.

In the immune system, dynamic cell-cell interactions coordinate bidirectional information transfer and control the potency and the scope of effector responses (*1*). One of the most important of these interactions is the immunological synapse (IS) formed between a cytotoxic T lymphocyte (CTL) and the infected or transformed target cell it aims to destroy (*2, 3*). IS formation is rapidly induced by recognition of cognate peptide-major histocompatibility complex (pMHC) on the target cell by T cell antigen receptors (TCRs) on the CTL. Once firm contact is established, the CTL secretes a toxic mixture of granzyme proteases and the hydrophobic protein perforin into the intercellular space. Perforin forms pores in the target cell membrane that stimulate the uptake of granzymes into the cytoplasm, where they induce apoptosis by cleaving specific substrates (*4*). Perforin and granzyme-mediated killing is the most prevalent mode of lymphocyte cytotoxicity, and it likely plays an important role in cellular immunotherapy approaches against cancer (*5*).

If and how the cytoskeletal architecture of the IS contributes to target cell killing remains an active area of study. IS formation is accompanied by dramatic reorganization of both microtubules and filamentous actin (F-actin) (*6*). Within minutes of TCR stimulation, the centrosome (also called the microtubule-organizing center) moves to a position just beneath the IS. The centrosome is closely associated with lytic granules, the secretory lysosomes that store perforin and granzyme, and its reorientation positions these granules next to the synaptic membrane (*3*). This promotes the directional secretion of granule contents into the intercellular space, enhancing both the potency and the specificity of target cell killing. The precise role of F-actin remodeling, by contrast, is less clear. Our current conception of synaptic F-actin is strongly influenced by imaging studies in which the target cell is replaced by a glass surface or a supported bilayer containing stimulatory pMHC or antibodies against the TCR. In this context, T cells form radially symmetric synapses characterized by intense F-actin accumulation at the periphery and depletion from the center (*7-10*). This annular configuration is thought to encourage lytic granule fusion at the center of the IS by clearing F-actin from the plasma membrane in this zone (*3, 11, 12*). Although this model is conceptually appealing, it is unclear how well it applies to granule release in bona fide CTL-target cell conjugates, where synaptic F-actin rings are less apparent and, when observed, often quite transient.

Synaptic F-actin is also highly dynamic, forming protrusions and lamellipodial sheets that exhibit both centripetal retrograde flow and radial anterograde movement (*10, 12-15*). Importantly, these dynamics enable T cells to impart mechanical force across the IS (*16, 17*). In CTLs, the capacity to exert synaptic force is strikingly correlated with cytotoxic potential (*18*). Biophysical and imaging experiments suggest that force enhances cytotoxicity by increasing the membrane tension of the target cell, which in turn promotes the pore forming activity of secreted perforin. How CTLs achieve this synergy between mechanical and secretory output, however, is not known.

Here, we applied microfabrication and high-resolution live imaging to investigate how CTLs mechanically potentiate the chemical activity of perforin, a process we refer to as mechanopotentiation. Using stimulatory micropillar arrays that trigger IS formation in three dimensions, we have found that lytic granule release occurs at the base of F-actin rich synaptic protrusions that extend into the antigen-presenting surface. These protrusions, which are generated by the Wiskott-Aldrich Syndrome protein (WASP) and the Arp2/3 actin nucleation complex, are required for synaptic force exertion and cytotoxic efficiency. Our results provide insight into how cells organize mechanical output and demonstrate how three-dimensional architecture influences the functionality of communicative interfaces in the immune system.

## RESULTS

### CTLs form synaptic protrusions on stimulatory micropillar arrays

Synaptic force exertion can be measured by imaging T cells on arrays of flexible polydimethylsiloxane (PDMS) micropillars bearing immobilized TCR ligands and adhesion proteins (Fig. 1A) (*17, 18*). T cells form IS-like contacts with these arrays and induce pillar deflections that can be converted into force vectors based on the known dimensions and composition of the pillars. This approach enables spatiotemporal correlation of mechanical output with other architectural components and cellular processes. In this manner, we previously found that lytic granule release (also called degranulation) tends to occur in regions of active pillar deflection (*18*). This result raised the possibility that there might be specific structures within the IS that mechanopotentiate perforin function by imparting force in close proximity to granule secretion.

**Figure 1.**
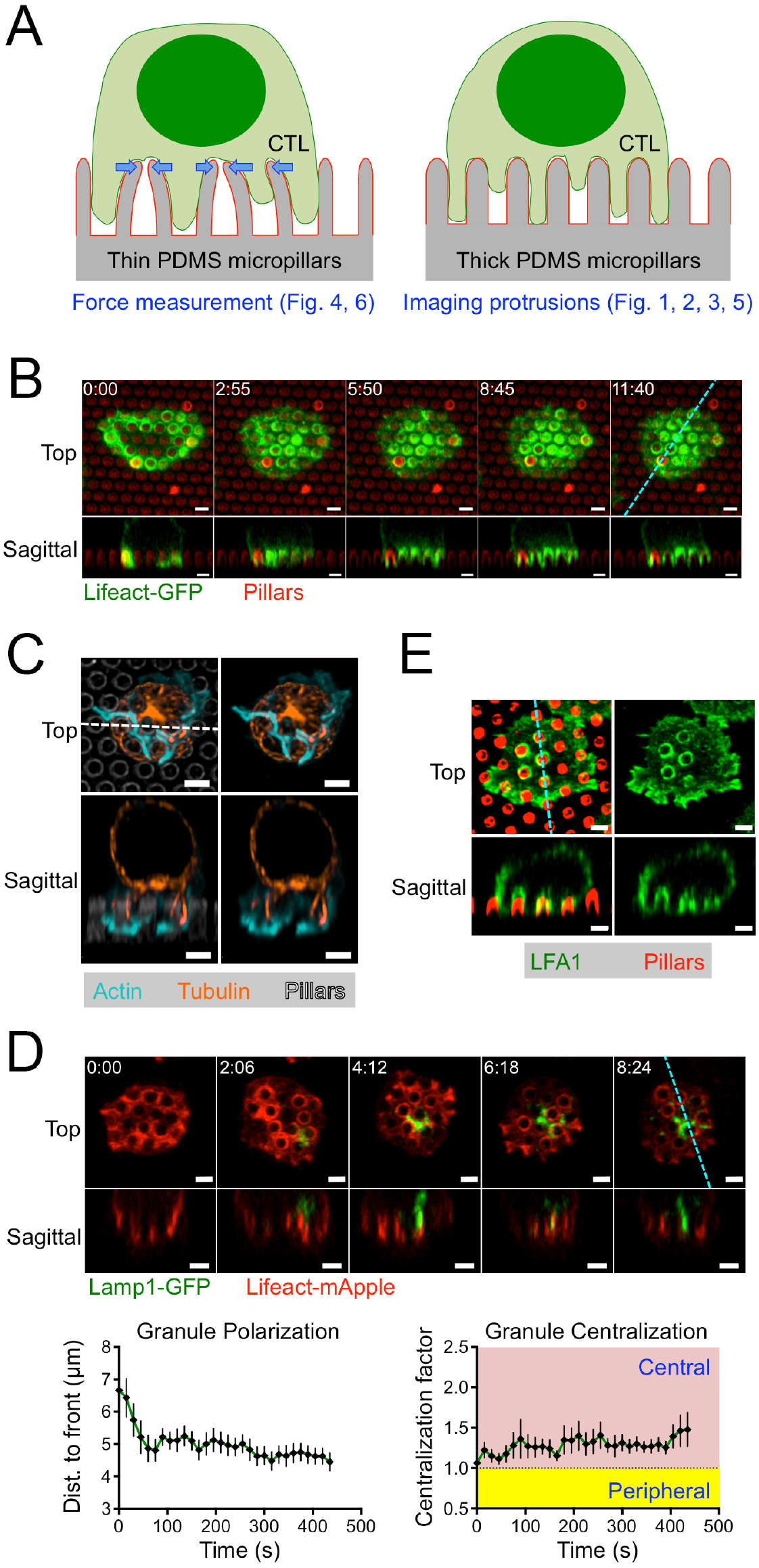
CTLs form protrusions on stimulatory PDMS micropillars. (A) Schematic diagram of tall, thin micropillars used for force measurements (left) and shorter, thicker micropillars used for imaging synaptic protrusions (right). (B-E) OT1 CTLs were imaged by confocal microscopy on micropillars bearing H2-K^b^-OVA and ICAM1. z-projection images (top views) are shown above with corresponding sagittal views below. Dotted lines (cyan in B, D, and E, white in C) denote the slicing plane used for the sagittal images in each panel. (B) Time-lapse montage of a representative CTL expressing Lifeact-GFP, with micropillars shown in red. (C) Fixed image of a representative CTL stained with phalloidin (to visualize F-actin) and anti-tubulin antibodies. Micropillars are shown in gray in the left images. (D) Above, time-lapse montage of a representative CTL expressing Lifeact-mApple and Lamp1-GFP. Below left, mean distance between the lytic granule cloud and the cell front, graphed against time. Below right, centralization factor analysis of Lamp1-GFP (see Materials and Methods). In both graphs, time 0 denotes initial contact with the pillars and error bars indicate standard error of the mean (SEM). N = 10. (E) Fixed image of a representative CTL stained with anti-LFA1 antibodies, with micropillars shown in red. All scale bars = 2 μm. In B and D, time in M:SS is indicated in the upper left corner of each top view image.

To identify candidate structures that could be involved in this mechanopotentiation, we closely examined the dynamic architecture of CTL synapses on micropillar arrays. For these experiments, we used primary CTLs expressing the OT1 TCR, which is specific for the ovalbumin_257-264_ peptide presented by the class I MHC protein H2-K^b^ (H2-K^b^-OVA). OT1 CTLs were retrovirally transduced with Lifeact-GFP, a fluorescent probe for F-actin, and imaged by confocal microscopy on micropillars coated with H2-K^b^-OVA and ICAM1, a ligand for the α_L_β_2_ integrin LFA1. The pillars in these arrays (1-1.5 μm in diameter and 4-5 μm tall) were thicker, shorter, and therefore more rigid than the pillars used for T cell force measurements (0.7 μm in diameter and 6 μm tall) (Fig. 1A). These thicker pillars are not substantially distorted by CTLs, and effectively function as a regularly crenulated stimulatory surface that facilitates quantitative assessment of IS growth in three dimensions.

Within minutes of initial contact with the arrays, the CTLs formed long, F-actin rich protrusions that invaded the spaces between adjacent pillars (Fig. 1B). Time-lapse experiments using both confocal and lattice light-sheet microscopy revealed that the F-actin in these protrusions was highly dynamic, coruscating up and down the length of each pillar (Fig. 1B, video S1). Periodically, F-actin free gaps appeared at the base of the protrusions, in the regions around the pillar tops. During protrusion growth, F-actin accumulation was often strongest at the leading edge, implying a causative relationship between actin polymerization and the formation of these structures (fig. S1). Most of the microtubule cytoskeleton, by contrast, was constrained to the region above the pillars, although individual microtubules were observed to extend into a subset of protrusions (Fig. 1C). In most cells, the centrosome reoriented to a position in the plane of the pillar tops, but did not proceed into the inter-pillar spaces. Hence, on micropillar arrays, CTLs form dynamic, F-actin rich protrusions at the IS that exclude the centrosome.

CTLs tended to elaborate synaptic protrusions in stages. During initial cell spreading, invasion into the micropillar zone was typically constrained to the periphery of the contact (Fig. 1B, video S1). However, once the radial size of the IS stabilized, after ~60 s, protrusions formed in the more central regions of the interface. The growth of these central protrusions was intriguing to us because prior studies had indicated that lytic granules accumulate in this portion of the IS (*3, 11, 19, 20*). To investigate the spatial relationship between synaptic protrusions and lytic granules more closely, we imaged CTLs expressing Lifeact-mApple together with a GFP-labeled form of the lysosomal protein Lamp1 (Lamp1-GFP). In most CTLs, lytic granules appeared as a cluster of distinct compartments within the cytoplasm. In the first two minutes of contact formation, this granule cluster moved downward, settling ~5 μm from the cell front, roughly at the level of the pillar tops (Fig. 1D). This reorientation behavior implied a close association between granules and the centrosome, as previously reported (*11, 20, 21*). Notably, after reorienting downward, the granules tended to occupy central, rather than peripheral, locations within the IS. We quantified this behavior by calculating the normalized proximity of granule fluorescence to the IS center of gravity (called the “centralization factor”) (fig. S2). This analysis revealed that the granules tended to be closer to the center of the IS than would be expected by chance (Fig. 1D).

The proximity of lytic granules to the base of synaptic protrusions at the center of the IS raised the possibility that these structures might be involved in cytotoxic mechanopotentiation after granule exocytosis. Indeed, synaptic protrusions were highly enriched in LFA1 (Fig. 1E), consistent with them being strongly adhesive and capable of exerting force. Previously, we found that cytolytic mechanopotentiation requires phosphoinositide 3-kinase (PI3K) signaling and is enhanced by the depletion of PTEN, a lipid phosphatase that antagonizes PI3K (*18*). shRNA-mediated suppression of PTEN augmented F-actin accumulation in synaptic protrusions (fig. S3), further supporting the idea that these structures transmit synaptic forces that promote cytotoxicity.

### Lytic granule release occurs at the base of synaptic protrusions

Degranulation can be detected in single cell imaging experiments with a fluorescent reporter containing a pH sensitive GFP (pHluorin) fused to the granule-targeting domain of Lamp1 (*22*). Within lytic granules, the low pH environment quenches the fluorescence of pHluorin-Lamp1. Granule fusion with the plasma membrane, however, neutralizes the pH around the reporter, leading to a rapid and localized increase in fluorescence. To explore the spatial relationship between degranulation and synaptic protrusions, OT1 CTLs expressing pHluorin-Lamp1 were imaged by confocal and lattice light-sheet microscopy on fluorescent micropillars coated with H2-K^b^-OVA and ICAM1 (Fig. 2A). Protrusions were visualized in these experiments either using Lifeact-mRuby2 or by staining with a fluorescent F_ab_ fragment against the surface marker CD45. After IS formation, degranulation events appeared as sudden flashes of GFP fluorescence, which were often visible for only one time point. Importantly, these events clustered close to the plane of the pillar tops (Fig. 2B-C, Video S2), the same vertical zone occupied by lytic granules and the centrosome. This position was well behind the leading edge of CTL protrusions, which extended approximately 5 μm into the inter-pillar space.

**Figure 2.**
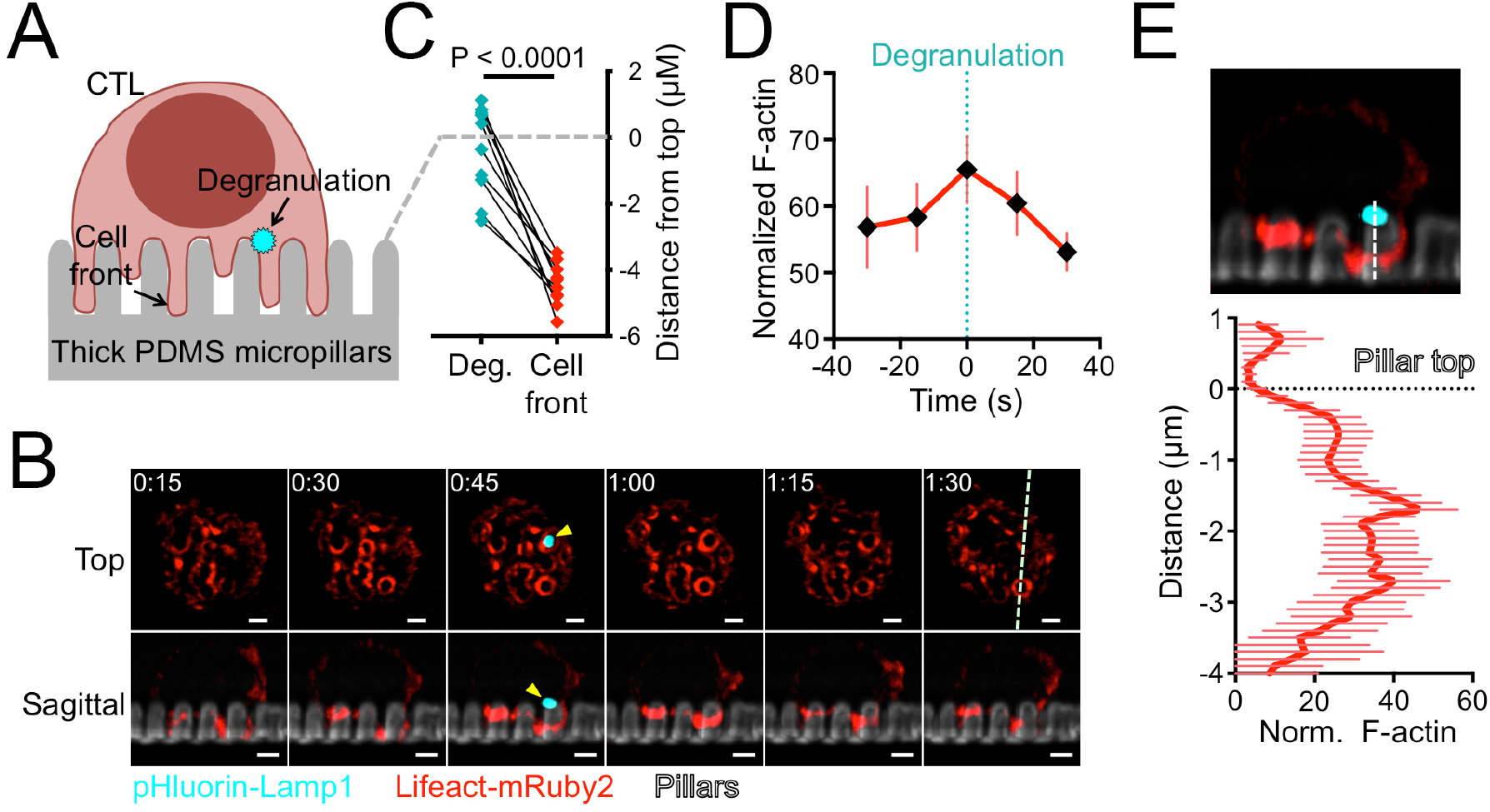
Degranulation occurs at the base of synaptic protrusions. (A) Schematic diagram showing CTL degranulation (visualized by pHluorin-Lamp1) on micropillar arrays. (B-E) OT1 CTLs expressing pHluorin-Lamp1 were imaged by confocal (C) or lattice light-sheet (B, D-E) microscopy on micropillars bearing H2-K^b^-OVA and ICAM1. (B) Time-lapse montage of a representative CTL expressing Lifeact-mRuby2 and pHluorin-Lamp1, with micropillars shown in gray. z-projection images (top views) are shown above with corresponding sagittal views below. The white dotted line denotes the slicing plane used for the sagittal images. Yellow arrowheads indicate the degranulation event. Time in M:SS is indicated in the upper left corner of each top view image. Scale bars = 2 μm. (C) Graph showing the vertical displacements of degranulation events (Deg., cyan) relative to the plane of the pillar tops (dotted gray line) along with the corresponding position of the cell front (red, visualized with a fluorescent F_ab_ fragment against CD45). N = 11 degranulations. P calculated from two-tailed paired Student’s T-test. (D) F-actin accumulation in the region of degranulation in z-projection images of CTLs expressing Lifeact-mRuby2 and pHluorin-Lamp1. Graph shows the average Lifeact-mRuby2 intensity within a 1 μm diameter circle centered on the degranulation position, starting two time points before the degranulation and ending two time points after. (E) Below, average Lifeact-mRuby2 intensity derived from linescans vertically bisecting the midpoint of the degranulation. The dotted black line denotes the plane of the pillar tops. Above, a representative image used for the analysis, with the linescan region indicated by the dotted white line. Error bars in D and E denote SEM. N = 18 degranulations.

Degranulation was not observed in large regions of sustained F-actin depletion. Rather, it appeared to be associated with the transient accumulation of F-actin in the zone directly beneath it (Fig. 2B, video S2). To quantify this effect, we calculated the normalized Lifeact-mRuby2 intensity in the immediate vicinity of each degranulation using z-projection images from multiple lattice light-sheet experiments. This analysis revealed an increase in local F-actin during granule release (Fig. 2D). Importantly, linescans of sagittal slice images demonstrated that F-actin did not overlap precisely with the granule release position. Rather, it tended to accumulate underneath, closer to the base of the pillars (Fig. 2E). We conclude that degranulation on micropillar arrays occurs in small F-actin free zones that form transiently at the base of active F-actin rich protrusions.

### Synaptic protrusions require the Arp2/3 complex

Having characterized the structure and dynamics of synaptic protrusions, we turned our attention to their molecular basis and biological function. We were particularly interested in the Arp2/3 complex, which nucleates actin polymerization from the sides of existing actin filaments (*23*). Previous studies had established Arp2/3 and its upstream regulators as critical drivers of synaptic F-actin remodeling (Fig. 3A)(*24-26*). To assess the importance of Arp2/3, we utilized CK666, a small molecule inhibitor of the complex (*27*). Treatment with CK666 dramatically attenuated protrusive activity on micropillar arrays. (Fig. 3B, video S3 and S4). We quantified these data by calculating the enrichment of F-actin accumulation (visualized using Lifeact-GFP) in the region beneath the pillar tops (fig. S4A). This analysis revealed a dose dependent reduction in synaptic protrusions (Fig. 3C). Notably, a small amount of protrusive activity was still observed even at high CK666 concentrations, possibly representing residual TCR-induced actin polymerization by formins (*24, 28*).

**Figure 3.**
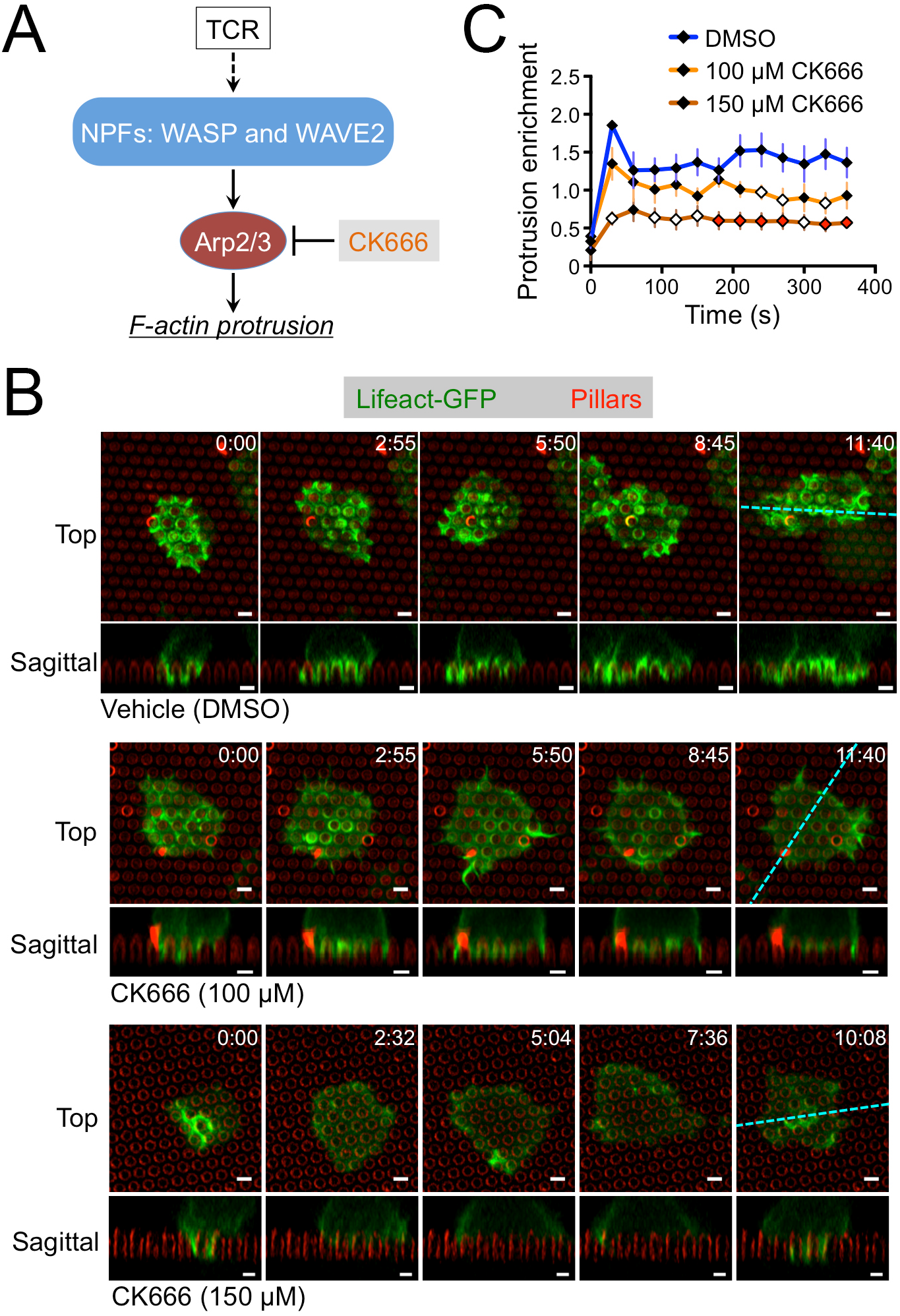
CK666 blocks protrusion formation. (A) Diagram showing TCR-induced activation of actin polymerization through NPFs and Arp2/3. (B-C) OT1 CTLs expressing Lifeact-GFP were imaged by confocal microscopy on fluorescent micropillars bearing H2-K^b^-OVA and ICAM1 in the presence of the indicated concentrations of CK666. (B) Time-lapse montages of representative CTLs, with micropillars shown in red. z-projection images (top views) are shown above with corresponding sagittal views below. Cyan dotted lines denote the slicing planes used for the sagittal images. Time in MM:SS is indicated in the upper right corner of each top view image. Scale bars = 2 μm. (C) Lifeact-GFP enrichment in protrusions was quantified over time as described in Materials and Methods, with time 0 indicating initial contact with the pillars. N = 5 cells for each condition, with error bars denoting SEM. Filled white diamonds and filled red diamonds indicate P < 0.05 and P < 0.01, respectively, calculated by two-tailed Student’s T-test comparing each CK666 condition against vehicle control.

The capacity of CK666 to block protrusion formation provided a strategy for determining the role of these structures in synaptic force exertion and cytotoxicity. To measure cellular forces, we imaged CTLs on stimulatory PDMS arrays containing narrow, deformable micropillars (Fig. 1A). Treatment with CK666 reduced force exertion by ~ 75% (Fig. 4A-B, videos S5 and S6), indicating that Arp2/3 dependent protrusive activity is indeed critical for IS mechanics. Next, we assessed the importance of Arp2/3 for cytotoxic function by incubating OT1 CTLs with OVA-loaded RMA-s target cells in the presence of CK666. Target cell lysis was inhibited by CK666 in a dose dependent manner (Fig. 4C), implying a critical role for Arp2/3 in the process. To better define the basis for this cytotoxicity defect, we examined several molecular and cellular events involved in T cell activation and target cell killing. CK666 treatment did not alter TCR-induced phosphorylation of Erk1/2 and Akt and had little effect on the degradation of IκB (fig. S4B). This suggested that signaling through the MAP kinase, PI3K, and NF-κB pathways was intact. By contrast, we observed a modest inhibition of TCR-induced calcium (Ca^2+^) flux in CK666 treated cells (fig. S4C-D). Elevated intracellular Ca^2+^ is known to be a prerequisite for lytic granule release (*19, 29*). Consistent with this idea, we found that CK666 significantly inhibited degranulation, which we measured by staining for surface exposed Lamp1 (Fig. 4D). We also observed a marked inhibition of CTL-target cell conjugate formation, as quantified by flow cytometry (Fig. 4E).

**Figure 4.**
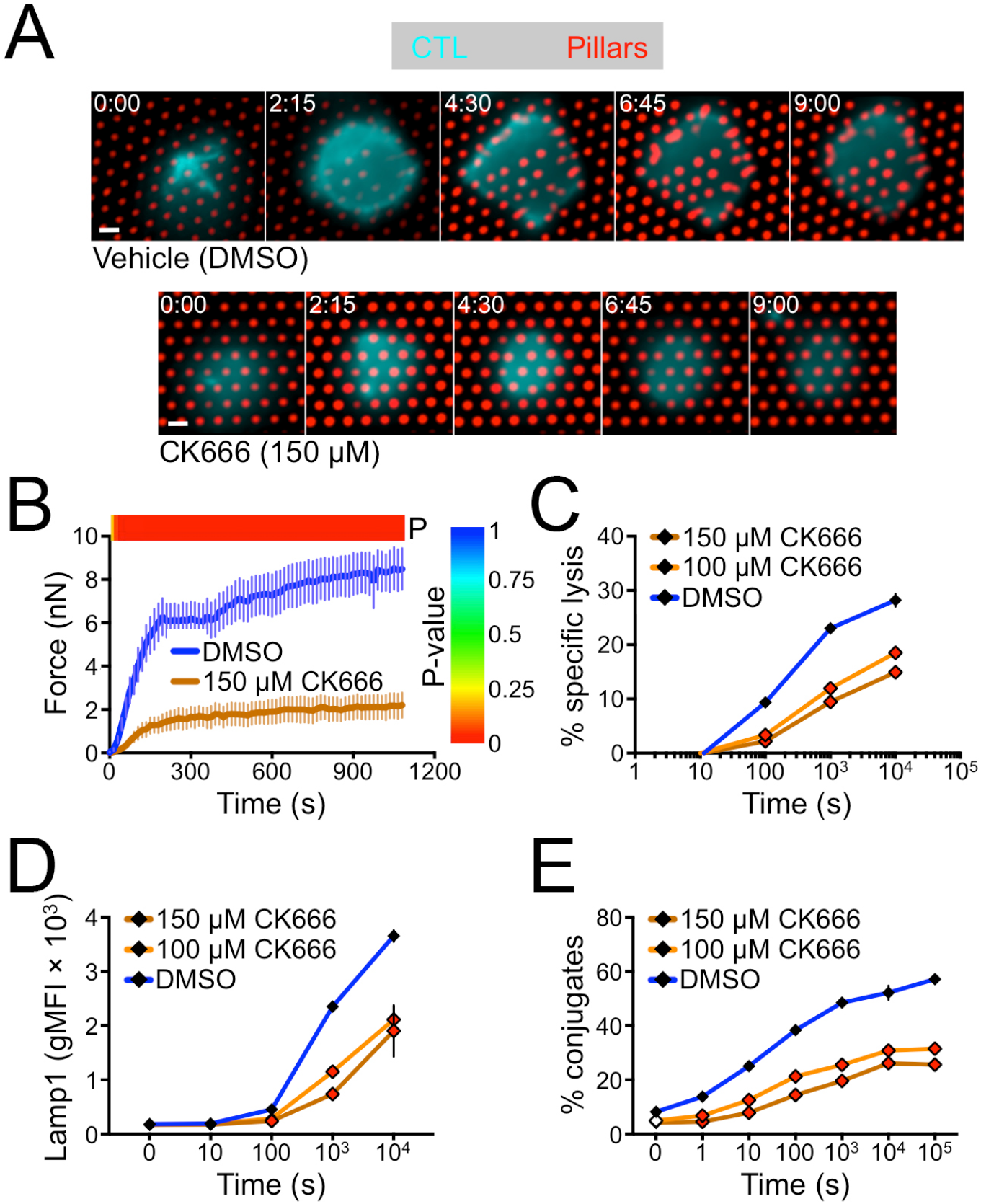
CK666 inhibits force exertion and cytotoxicity. (A-B) OT1 CTLs were labeled with a fluorescent anti-CD45 F_ab_, incubated with 150 μm CK666 or vehicle control (DMSO) as indicated, and imaged on narrow fluorescent micropillars coated with H2-K^b^-OVA and ICAM1. (A) Time-lapse montages of representative CTLs showing pillar deflection. Time in M:SS is indicated in the upper left corner of each image. Scale bars = 2 μm. (B) Total force exertion against pillar arrays was graphed versus time. Color bar above the graph indicates the P-value for each time point (two-tailed Student’s T-test). N = 9 for DMSO, 10 for CK666. (C-E) RMA-s target cells were loaded with increasing concentrations of OVA and then mixed with CTLs in the presence or absence of CK666 as indicated. (C) Specific lysis of RMA-s cells. (D) Degranulation measured by surface exposure of Lamp1. (E) CTL-target cell conjugate formation measured by flow cytometry. All error bars denote SEM. In C-E, filled white diamonds and filled red diamonds indicate P < 0.05 and P < 0.01, respectively, calculated by two-tailed Student’s T-test comparing each CK666 condition against the DMSO control.

Taken together, these data demonstrated that the Arp2/3 complex is required for the formation of synaptic protrusions and also for several TCR dependent responses associated with cytotoxicity: synaptic force exertion, degranulation, and strong adhesion to the target cell. This pattern of results was not inconsistent with our hypothesis that synaptic protrusions enhance killing via cytolytic mechanopotentiation. We could not, however, exclude the possibility that the killing defect induced by CK666 was caused entirely by reduced CTL degranulation and/or conjugate formation, either of which may have occurred independently of the protrusion defect. Nor could we rule out the possibility that the cytotoxicity phenotype resulted from effects of CK666 on the target cells. Hence, in order to define the role of synaptic protrusions unambiguously, it was necessary to establish more specific molecular perturbations.

### WASP and WAVE2 control spatially and functionally distinct synaptic protrusions

The localization and activity of the Arp2/3 complex is controlled by the nucleation promoting factor (NPF) family of actin regulators (*23*). Among NPFs, both WASP and WAVE2 have been implicated in synaptic F-actin remodeling (Fig. 3A). WAVE2, which is activated by the GTP-bound form of the small GTPase Rac, is thought to promote cell spreading and adhesion during IS formation (*30-32*). WASP, for its part, functions downstream of the GTPase Cdc42 and the adaptor protein Nck, and it has been linked to IS stability and the formation of protrusive structures during diapedesis and antigen scanning (*25, 33-35*). To begin investigating the role of each protein in the formation of synaptic protrusions, we imaged OT1 CTLs expressing GFP-labeled forms of WASP and WAVE2 on stimulatory micropillars. WAVE2-GFP accumulated strongly in the periphery of the IS during initial cell spreading (< 1 min, Fig. 5A, video S7). In subsequent time points, transient bursts of WAVE2-GFP regularly appeared in isolated peripheral domains (Fig. 5A, magenta arrowheads) These lateral bursts of WAVE2-GFP often occurred concomitantly with movement of the IS toward the same side (video S7). By contrast, WASP-GFP accumulated in annular structures that encircled individual pillars in central and intermediate synaptic domains (Fig. 5A, video S8). WASP-GFP exhibited a significantly higher mean centralization factor than WAVE2-GFP at all time points (Fig. 5B, fig. S2), confirming that WASP adopts a more centralized localization pattern than WAVE2. Collectively, these data indicated that WAVE2 accumulates within peripheral structures involved in lateral motion, while WASP associates with protrusions closer to the center of the IS.

**Figure 5.**
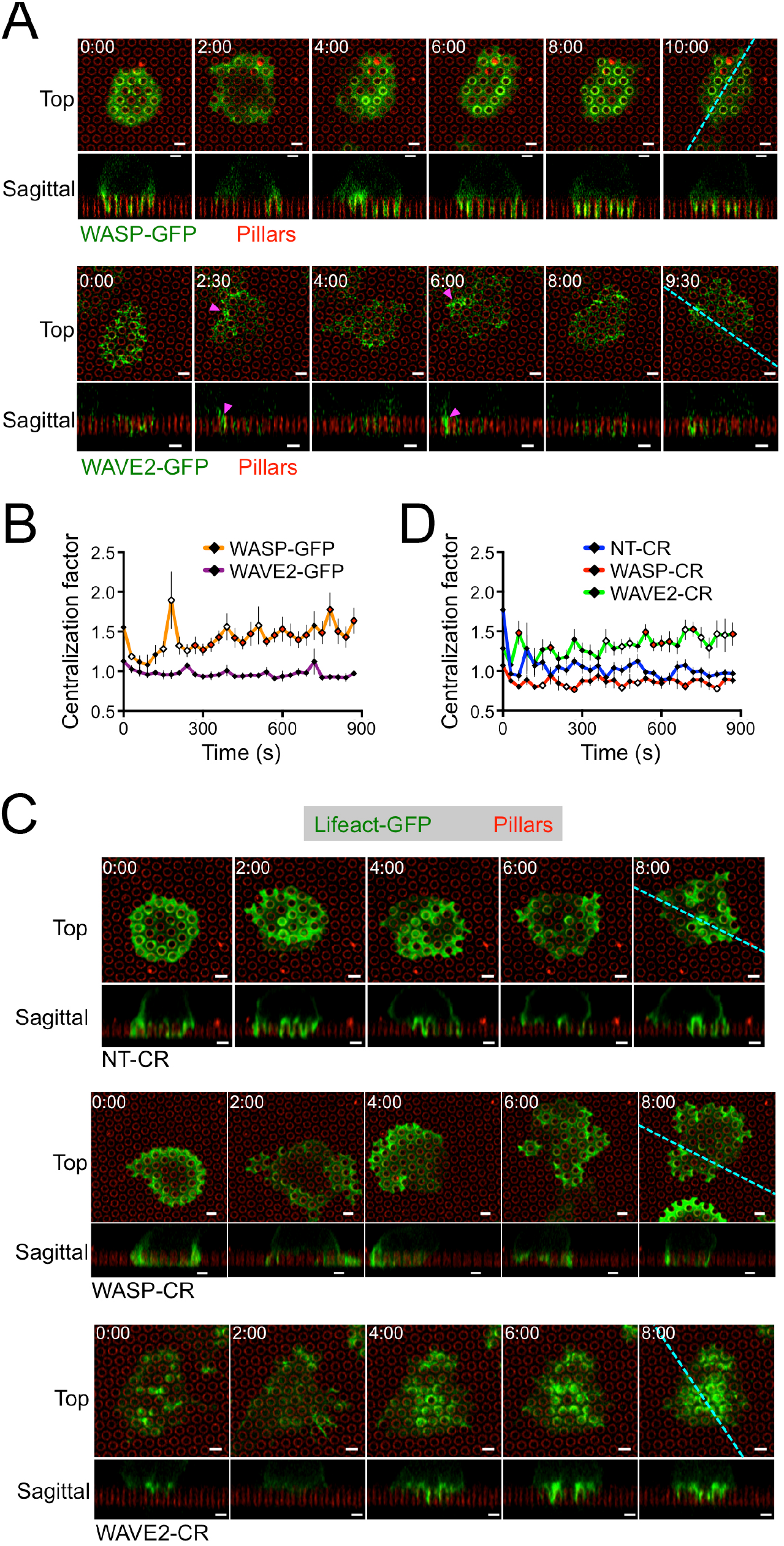
WASP and WAVE2 control distinct subsets of protrusions. (A-B) OT1 CTLs expressing WASP-GFP or WAVE2-GFP as indicated were imaged by confocal microscopy on fluorescent micropillars bearing H2-K^b^-OVA and ICAM1. (A) Time-lapse montages of representative CTLs, with micropillars shown in red. z-projection images (top views) are shown above with corresponding sagittal views below. Cyan dotted lines denote the slicing planes used for the sagittal images. Magenta arrowheads indicate representative lateral accumulations of WAVE2-GFP. (B) Centralization factor analysis of WASP-GFP and WAVE2-GFP (see Materials and Methods), with time 0 denoting initial contact with the pillars. N = 6 for each cell type. (C-D) NT-CR, WASP-CR, and WAVE2-CR CTLs expressing Lifeact-GFP were imaged by confocal microscopy on fluorescent micropillars bearing H2-K^b^-OVA and ICAM1. (C) Time-lapse montages of representative CTLs, with micropillars shown in red. z-projection images (top views) are shown above with corresponding sagittal views below. Cyan dotted lines denote the slicing planes used for the sagittal images. (D) Centralization factor analysis of Lifeact-GFP in NT-CR, WASP-CR, and WAVE2-CR OT1 CTLs, with time 0 denoting initial contact with the pillars. N = 6 for each cell type. In all montages, time in MM:SS is indicated in the upper left corner of each top view image. Scale bars = 2 μm. In graphs, filled white diamonds and filled red diamonds indicate P < 0.05 and P < 0.01, respectively, calculated by two-tailed Student’s T-test comparing WASP-GFP to WAVE2-GFP (B) or WASP-CR and WAVE2-CR to NT-CR (C). Error bars denote SEM.

To assess the importance of WASP and WAVE2 for IS remodeling in three dimensions, we developed a CRISPR/Cas9 approach to target the *Was* and *Wasf2* genes, respectively. Using optimized gRNAs, we were able to achieve highly efficient protein depletion in OT1 CTLs (fig. S5A and Materials and Methods). WASP or WAVE2 deficient CTLs prepared in this manner (WASP-CR and WAVE2-CR, respectively) were transduced with Lifeact-GFP and then imaged on micropillar arrays to visualize synaptic protrusions. Whereas control CTLs expressing nontargeting gRNA (NT-CR) formed protrusions in both the center and the periphery of the IS, the protrusive activity of WASP-CR cells was largely constrained to the periphery (Fig. 5C, videos S9 and S10). WASP-CR synapses also tended to be more laterally mobile, consistent with previous work (*33*). In WAVE2-CR CTLs, by contrast, protrusions appeared to concentrate in the central domain (Fig. 5C, Video S11). Centralization factor analysis over the entire data set revealed that the F-actin distributions of WAVE2-CR CTLs were more centralized than those of NT-CR controls, which were in turn more centralized than those of WASP-CR CTLs (Fig. 5D). Hence, WASP deficiency leads to a specific loss of central protrusions, whereas WAVE2 deficiency eliminates peripheral structures. These results, which mirror the localization patterns of WASP-GFP and WAVE2-GFP, indicate that WASP and WAVE2 control spatially distinct subsets of synaptic protrusions.

Next, we investigated the mechanical consequences of WASP and WAVE2 depletion by imaging NT-CR, WASP-CR, and WAVE2-CR CTLs on narrow micropillar arrays (Fig. 1A and 6A, videos S12-S14). WASP-CR CTLs avidly engaged the arrays and began to deform them as quickly as did NT-CR controls. The overall magnitude of WASP-CR force exertion, however, was significantly reduced (Fig. 6B). By contrast, depletion of WAVE2 delayed the onset of force exertion but did not affect its overall magnitude (Fig. 6A and 6B). To assess the spatial patterns of these mechanical responses, we plotted the number of strongly deflected pillars as a function of radial distance from the IS COG (fig. S5B). NT-CR and WAVE2-CR CTLs induced pillar deflections in both the central IS (< 3 μm from the COG) and in the periphery (> 3 μm from the COG). By contrast, in WASP-CR CTLs there was a marked absence of centrally localized events (Fig. 6C). Hence, the capacity to generate protrusions in the center of the IS was associated with force exertion in that domain.

**Figure 6.**
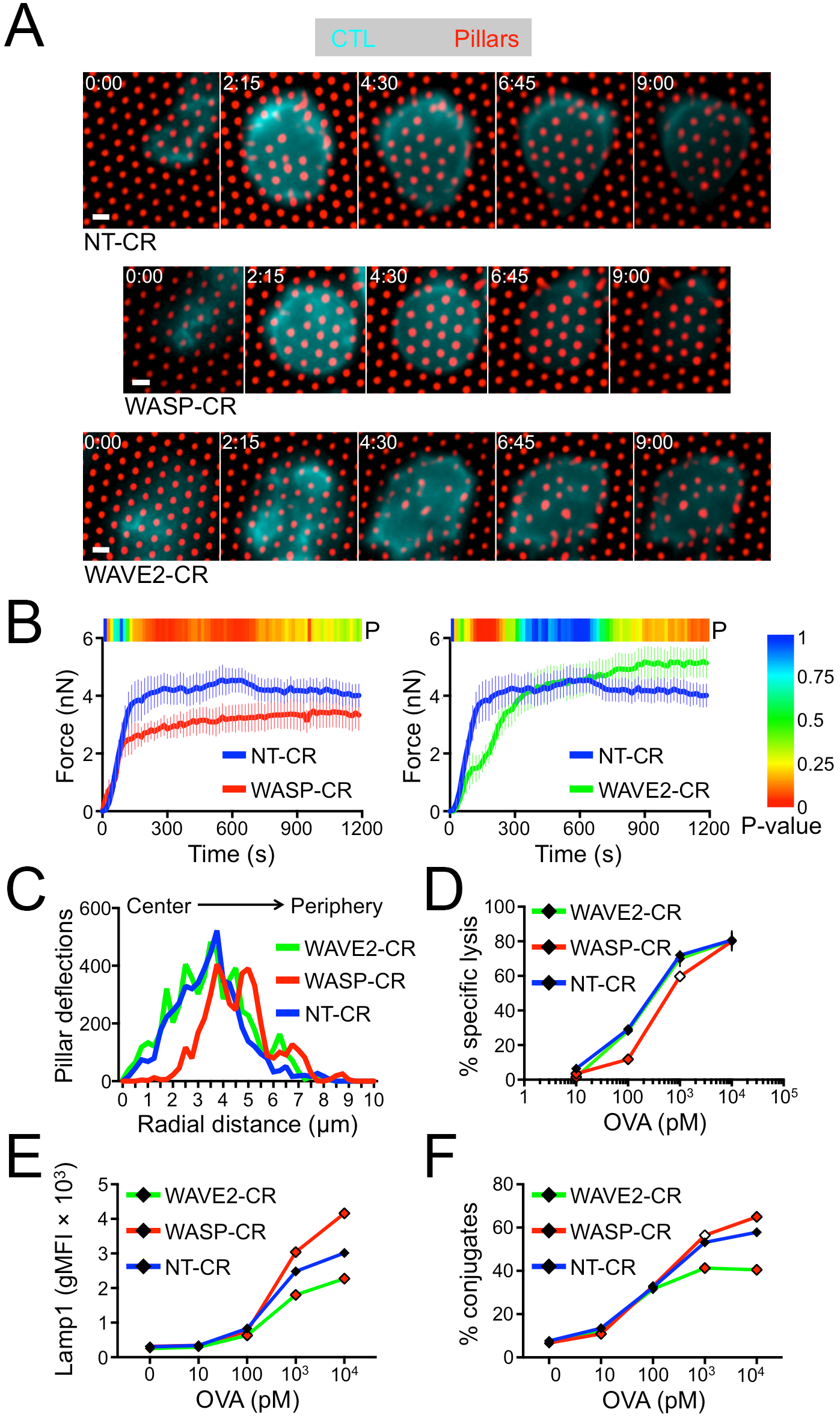
WASP and WAVE2 depletion induce distinct functional phenotypes. (A-B) NT-CR, WASP-CR, and WAVE2-CR OT1 CTLs were labeled with a fluorescent anti-CD45 F_ab_ and imaged on narrow fluorescent micropillars coated with H2-K^b^-OVA and ICAM1. (A) Time-lapse montages of representative CTLs showing pillar deflection. Time in M:SS is indicated in the upper left corner of each top view image. Scale bars = 2 μm. (B) Total force exertion against pillar arrays was graphed versus time. Color bar above each graph indicates the P-value for each time point (two-tailed Student’s T-test). (C) Histogram showing the distribution of strong deflections as a function of radial distance from the center of the IS. N = 10 for each cell type in B and C. (D-F) RMA-s target cells were loaded with increasing concentrations of OVA and then mixed with NT-CR, WASP-CR, or WAVE2-CR CTLs. (D) Specific lysis of RMA-s cells. (E) Degranulation measured by surface exposure of Lamp1. (F) CTL-target cell conjugate formation measured by flow cytometry. All error bars denote SEM. In D-F, filled white diamonds and filled red diamonds indicate P < 0.05 and P < 0.01, respectively, calculated by two-tailed Student’s T-test comparing WASP-CR and WAVE2-CR to NT-CR.

Finally, we examined the cytotoxic function of CTLs lacking WASP and WAVE2. Depletion of WASP induced a significant defect in target cell killing (Fig. 6D), which was most pronounced (up to 50% reduction) at low levels of stimulatory OVA peptide. At higher OVA concentrations, however, killing by NT-CR and WASP-CR CTLs was equivalent. This killing phenotype was not associated with lower levels of degranulation (Fig. 6E), indicating that the reduced cytotoxicity of WASP-CR CTLs could not be attributed to a defect in granule release. Depletion of WAVE2 led to a distinct and somewhat variable cytotoxicity phenotype. In some experiments, we found little to no change in killing and degranulation, while in others we observed modest reductions that were most pronounced at high OVA concentrations (Fig. 6D, 6E, fig. S5C). Interestingly, WAVE2-CR CTLs, but not their WASP-CR counterparts, exhibited significantly reduced conjugate formation (Fig. 6F), implying that WAVE2 plays the more important role in target cell adhesion. Consistent with this interpretation, depletion of WAVE2, but not WASP, impaired CTL adhesion to ICAM1-coated surfaces, both in the presence and the absence of H2-K^b^-OVA (fig. S5D). We also examined indices of TCR signaling, and found that WASP-CR and WAVE2-CR CTLs exhibited normal TCR-induced Ca^2+^ flux and signaling through the MAPK, PI3K, and NF-κB pathways (fig. S5E-F). Hence, depletion of WASP or WAVE2 does not broadly disrupt early T cell activation.

We conclude that WASP plays a more important role than WAVE2 in boosting cytotoxicity and that it does so in a manner independent of TCR signaling, conjugate formation, and granule release. In interpreting the WASP-CR phenotype, it is interesting to note that the killing defect was strongest at low OVA concentrations, when degranulation was weaker and perforin levels limiting, and that it disappeared at high OVA concentrations, when perforin was abundant. Previous studies of human CTLs lacking WASP revealed a similar cytotoxicity phenotype: reduced killing (despite normal conjugate formation and degranulation), which was rescued by strong TCR stimulation (*34, 36*). This is precisely the pattern of results one would expect after blocking a mechanical process that boosts the per molecule efficiency of perforin. Taken together with the imaging data described above, these data suggest a model in which centralized, WASP dependent protrusions enhance target cell killing through cytolytic mechanopotentiation.

### WASP dependent protrusions distort target cells at the immunological synapse

If synaptic protrusions mechanopotentiate perforin function, then they should be capable of physically manipulating the target cell surface. To investigate this hypothesis, we performed live imaging experiments using H2-K^b+^ murine endothelial cells as targets for OT1 CTLs. In culture, endothelial cells adopt a flat, stellate architecture that is stable over time. Hence, deviations in this morphology within the IS can be attributed to the physical activity of the T cell (Fig. 7A). Indeed, previous studies have used endothelial cells in this manner to visualize shape changes induced by T cells (*13, 25*).

**Figure 7.**
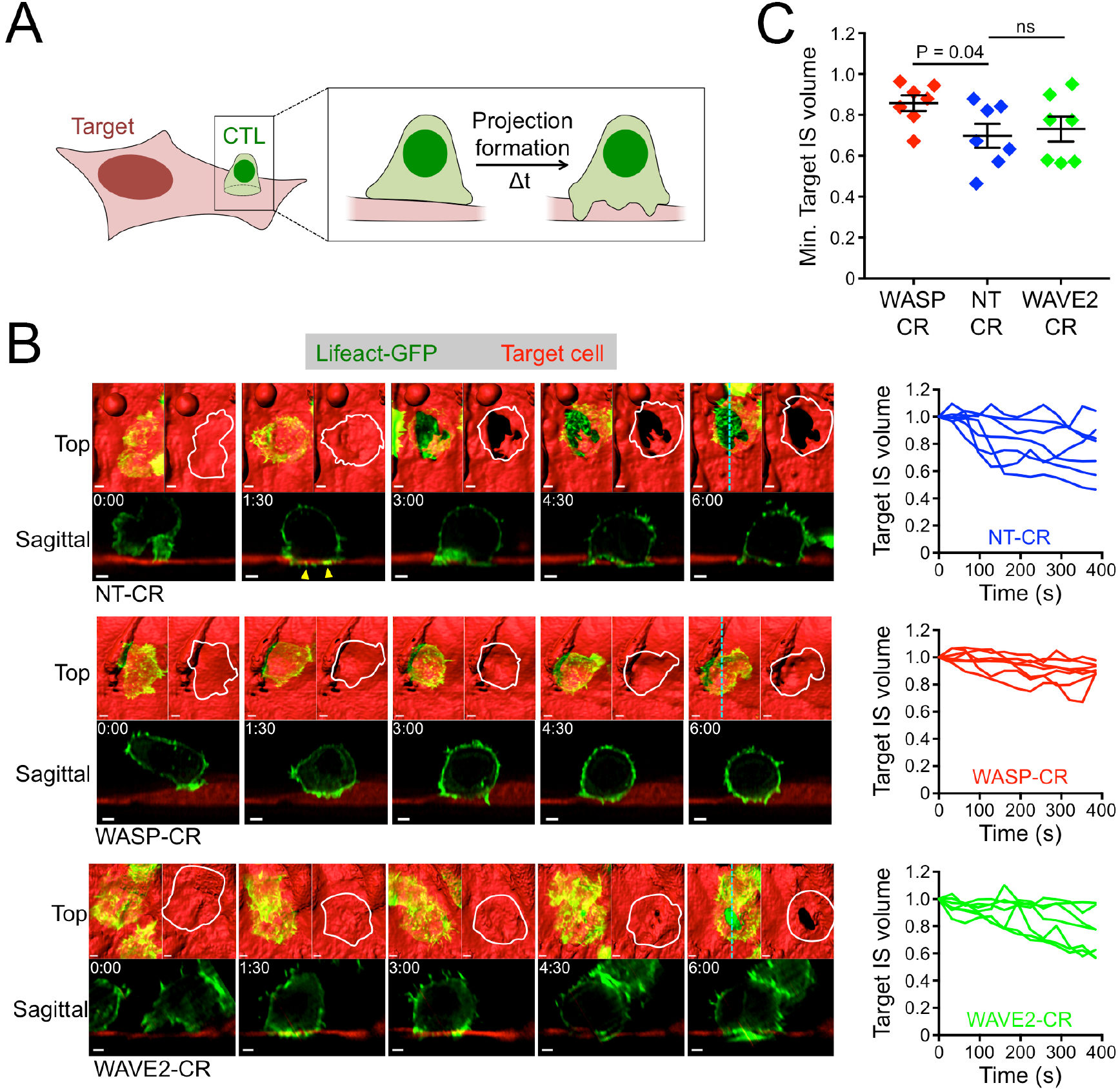
WASP drives target cell distortion at the IS. (A) Schematic diagram of a CTL inducing distortion of an adherent target cell. (B-C) NT-CR, WASP-CR, and WAVE2-CR OT1 CTLs expressing Lifeact-GFP were applied to confluent cultures of OVA-loaded endothelial target cells expressing iRFP670 and then imaged using lattice light-sheet microscopy. (B) Left, time-lapse montages of representative “vertically” oriented synapses, with z-projection images (top views) shown above and corresponding sagittal views below. Cyan dotted lines denote the slicing planes used for the sagittal images. In z-projection images, target cells are visualized by surface representation. Two z-projections are shown for each time point; Lifeact-GFP is shown on the left and the outline of the CTL of interest on the right. Time in M:SS is indicated in the upper left corner of each sagittal image. Scale bars = 2 μm. Yellow arrowheads denote protrusive structures in the NT-CR CTL that invade the target cell space. Right, target IS volume (see Materials and Methods) graphed against time, with time 0 denoting IS initiation. Each line corresponds to one CTL-target cell conjugate. (C) Graph of minimum target IS volume values achieved during the first 400 s of conjugate formation. N = 7 for each cell type. Error bars denote SEM. P calculated by two-tailed Student’s T-test.

To facilitate the imaging of cellular volume, we prepared endothelial cell lines that expressed mApple or iRFP670 uniformly in both the cytoplasm and the nucleus. These target cells were loaded with OVA, mixed with OT1 CTLs expressing fluorescently labeled Lifeact, and imaged by lattice light-sheet microscopy. Synapses formed readily and could be identified by their stability as well as the strong accumulation of interfacial F-actin within the CTL. Within minutes of IS initiation, CTLs generated small, protrusive F-actin structures that invaded the space occupied by the target cell (Fig. 7B, yellow arrowheads). This was followed shortly thereafter by rapid effacing of the target surface, which was most obvious in conjugates where the CTL attacked the target from above (Fig. 7B, video S15). In these vertically oriented synapses, the CTLs often appeared to displace the target cell completely. This displacement typically occurred before any obvious signs of target cell blebbing, suggesting that it was not part of the apoptotic cascade. Indeed, CTLs lacking perforin also formed large holes in target cells (fig. S6A), further supporting the idea that synaptic distortions result from a physical, rather than a chemical, process. TCR engagement was critical for these mechanical effects. In the absence of antigen, both the speed and the magnitude of distortion diminished substantially (fig. S6B), consistent with previous work (*13*). Finally, imaging of CTLs expressing Lamp1-GFP revealed that lytic granules accumulated close to areas of active target cell displacement (fig. S6C), implying that physical manipulation of the target cell contributes to perforin-and granzyme-mediated killing.

Next, we investigated the molecular basis of target cell distortion by comparing synapses formed by NT-CR, WASP-CR, and WAVE2-CR CTLs. Depletion of WASP markedly inhibited physical manipulation of the target surface. Although WASP-CR CTLs still exhibited F-actin accumulation at the IS, the displacement of target cell volume was less pronounced and more transient (Fig. 7B, video S16). To quantify this effect, we determined the volume beneath the CTL occupied by the target cell at a given time point and normalized this value to the volume occupied by the target cell in that same region before IS formation (fig. S6D). Analysis of this “Target IS volume” parameter confirmed that WASP depletion significantly reduced the distortive capacity of CTLs (Fig. 7B-C). CTLs lacking WAVE2 exhibited a qualitatively distinct phenotype; although they were still capable of substantial target cell distortion, their mechanical responses were somewhat delayed relative to those of NT-CR controls (Fig. 7B-C, video S17). Collectively, these results mirror the force exertion analysis of WASP-CR and WAVE2-CR cells (Fig. 6A-C), and they suggest that WASP dependent synaptic protrusions play a particularly important role in the physical distortion of target cells.

## DISCUSSION

It has been difficult to relate the architectural features of cell-cell interactions to their communicative functions because there are few experimental approaches for visualizing and perturbing small, dynamic cellular structures. The cytotoxic IS, a specialized interface that instructs target cells to die, boosts perforin toxicity by coordinating its secretion with the exertion of mechanical force. In the present study, we found that perforin release occurs at the base of dynamic F-actin rich protrusions that depend on WASP for their formation (Fig. 8). These WASP dependent protrusions were necessary for synaptic force exertion, particularly in more central regions of the IS close to lytic granules. They were also required for physical distortion of target cells in bona fide cytolytic interactions. Importantly, WASP deficient CTLs exhibited a defect in killing that could not be explained by reduced degranulation or conjugate formation. Taken together, these data identify synaptic protrusions as key components of a physical delivery system that enables CTLs to kill target cells with high efficiency. In this manner, CTLs are quite similar to venomous snakes and scorpions, which also use physical delivery systems to maximize the power of toxic secretions.

**Figure 8.**
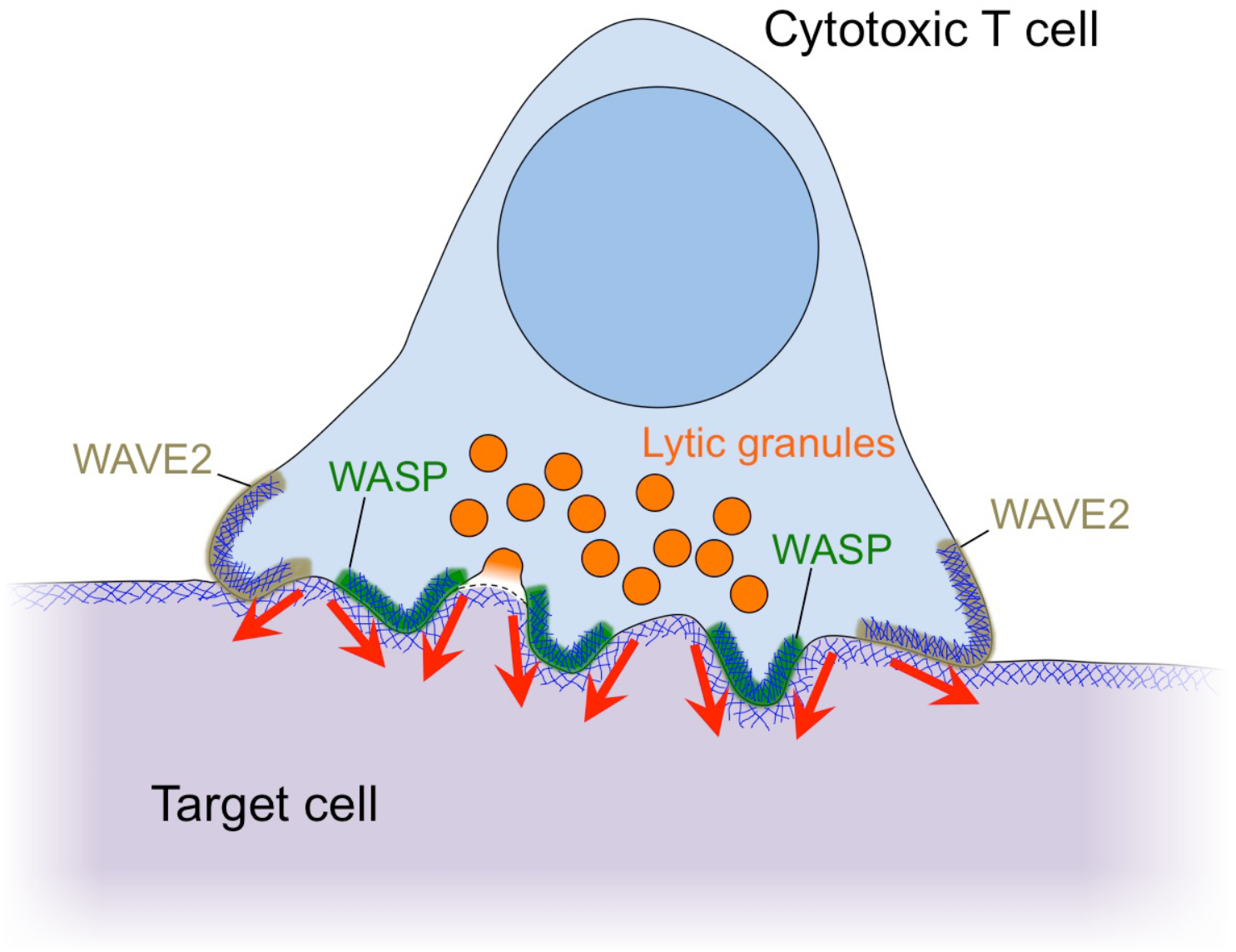
Cytolytic mechanopotentiation by WASP-dependent synaptic protrusions. A schematic diagram of the cytolytic IS showing peripheral WAVE2-dependent protrusions and central WASP-dependent protrusions. Red arrows denote force exertion.

In putting forth this model, we do not suggest that synaptic protrusions are a prerequisite for lytic granule release. Indeed, multiple groups have demonstrated that rigid stimulatory surfaces induce robust degranulation in the absence of centralized protrusive activity (*19, 22, 37-39*). That being said, it is tempting to consider the converse relationship, namely that lytic granule docking and fusion might modulate the architecture and mechanical activity of nearby protrusions. Consistent with this idea, we have documented a temporal correlation between degranulation events and increased F-actin accumulation in nearby protrusion tips. This observation raises the intriguing possibility that granules themselves could trigger the mechanopotentiation of their own secreted perforin.

The marked concentration of degranulation events at the base of synaptic protrusions implies that mechanisms exist for granule targeting to these domains. Previous studies have highlighted the importance of F-actin clearance for enabling granule access to the plasma membrane (*11, 12, 22, 39*). This is generally consistent with our observation that, on micropillar arrays, degranulation occurs in transient F-actin free regions at the base of synaptic protrusions. However, the presence of other F-actin hypodense areas in the CTL, which are not targeted by granules, implies that other factors contribute to the process. Lipid second messengers are known to influence exocytosis and membrane trafficking in a variety of cellular contexts (*40*). Among these, diacylglycerol is a particularly interesting candidate because it tends to accumulate in central synaptic domains that experience F-actin depletion (*41-43*). Granule delivery via microtubules is another intriguing possibility. Microtubules play a well-established role in trafficking granules from the back of the CTL toward the centrosome (*3*). Whether a different set of microtubules then guides the granules from the centrosome toward fusion sites in the synaptic membrane, however, is more controversial (*11, 44, 45*). We have observed that a small subset of microtubules extends into synaptic protrusions, and it will be interesting to explore if and how this subset contributes to granule exocytosis. Finally, it is possible that WASP itself plays a role. Indeed, a recent super-resolution imaging study demonstrated that WASP promotes degranulation close to regions of integrin clustering (*46*), implying a role for the protein in granule targeting.

T cells form a variety of protrusive structures at both interfacial and noninterfacial surfaces, which have been documented previously by electron microscopy and high-resolution fluorescence imaging (*13, 14, 20, 35, 47, 48*). Although the spatial distribution of these protrusions and their dynamics have implied roles in antigen scanning, signaling, and motility, their precise functions have in most cases remained undefined. Probably the best studied of these structures are the invadosome-like protrusions (ILPs), which were first observed in T cells initiating diapedesis through endothelial monolayers (*35*). Subsequently, ILPs were also found in antigen-induced synapses formed between T cells and endothelial cells, dendritic cells, or B cells (*13*). ILPs are podosomal structures that are enriched in LFA1 and require WASP and Arp2/3 for their formation (*25, 35*). The synaptic protrusions we have observed share these characteristics, and it is therefore tempting to speculate that they are a form of ILP. That CTLs might use the same cellular structure to facilitate both diapedesis and target cell killing highlights an underappreciated similarity between the two processes. Both rely on the physical manipulation of other cells through direct contact, in the first case to facilitate transmigration and in the second to promote destruction.

Synaptic ILPs have been observed to colocalize with the TCR and several membrane proximal signaling proteins, implying that ILPs promote TCR signaling (*13*). Consistent with this idea, WASP deficient T cells display defects in TCR-induced activation of phospholipase C-γ and Ca^2+^ flux under certain conditions (*25, 34, 49, 50*). These signaling defects are subtle, however, and it is unclear whether they fully explain the cytotoxicity phenotype that we and others have observed (*34, 36*). Indeed, our data indicate that WASP deficient CTLs kill poorly despite normal TCR signaling, Ca^2+^ flux, and degranulation, revealing a distinct role for WASP during the effector phase of the cytolytic response. This raises the possibility that WASP dependent protrusions may contribute to specific effector pathways in other immune cell-cell interactions. WASP is expressed by most cells of hematopoietic lineage and has been implicated in a variety of activation-induced events, including phagocytosis, receptor downregulation, and cytokine secretion (*51*). It will be interesting to investigate the extent to which its protrusive activity contributes to these processes.

Although this is the first instance, to our knowledge, of F-actin-rich protrusions being implicated in cellular cytotoxicity, studies in myoblast fusion have established a role for structures like these in modulating membrane integrity. Myoblast fusion is initiated by the formation of a WASP dependent protrusion on the “attacking” myoblast (*52*). This podosome-like structure (PLS), which is actually a cluster of small protrusions, promotes close apposition of cell membranes and has been observed to distort the surface of the opposing cell (*53*). Interestingly, myosin dependent tensioning of the opposing cell has been shown to facilitate the fusion process by countering the forces generated by the PLS (*54*). Similarly, we have found that enhanced tension in the target cell promotes cytotoxicity by accelerating perforin pore formation (*18*). The parallels between these two systems suggest that force exertion by WASP dependent protrusions may serve as a general mechanism for the destabilization of target membranes.

In marked contrast to WASP, WAVE2 accumulated in peripheral synaptic protrusions, and CTLs lacking WAVE2 exhibited a marked defect in adhesion to ICAM1-coated surfaces and conjugation with target cells. These observations suggest a role in cell spreading and adhesion, which is consistent with previous reports (*30-32*). Interestingly, WAVE2 depletion only weakly affected synaptic force exertion and killing, indicating that the protein and the peripheral structures it generates are not involved in cytolytic mechanopotentiation. These results do not exclude the possibility that WAVE2 might promote cytotoxicity in other settings, particularly in circumstances where target cells are more limiting and robust migration and adhesion are required for their destruction. What is clear from our data, however, is that the functionality of synaptic protrusions is partitioned both spatially (center versus periphery) and molecularly (WASP versus WAVE2) within the IS.

In vitro systems that collapse the IS into two dimensions have been invaluable for investigating its structure and function (*55, 56*). The inherent constraints of these systems, however, have limited our understanding in significant ways. Indeed, it was only by analyzing the IS in an oriented, three-dimensional environment that we were able to assess, in a controlled manner, the formation of synaptic protrusions and to study the implications of these structures for IS mechanics and effector responses. Micropatterned reconstitution systems like the ones employed here can reveal unexplored facets of interfacial architecture, and we anticipate that they will become increasingly important in future studies of complex cellular behavior.

## MATERIALS AND METHODS

### Study Design

The goal of this study was to understand how CTLs combine cytolytic secretion with force exertion at the IS. To this end, we employed fluorescence imaging of mouse CTLs, single cell biophysical assays, and functional experiments. Micropatterned PDMS substrates were used both to visualize the growth of synaptic protrusions and also to measure mechanical activity at the IS. To perturb protrusion formation and synaptic F-actin dynamics, we employed CRISPR/Cas9 technology, shRNA, and the Arp2/3 inhibitor CK666. Experimental sample sizes were not predetermined, and there were no predefined study endpoints. Experiments were not randomized, and investigators were not blinded during acquisition and data analysis. In general, experiments were performed at least twice (two biological replicates). Specific information about analysis methods can be found in the Image Analysis section below.

### Constructs

Retroviral expression constructs for Lifeact-GFP, Lifeact-mRuby2, Dyn1PH-GFP, and pHluorin-Lamp1 were previously described (*8, 22*). The Lamp1-GFP, WASP-GFP, and WAVE2-GFP constructs were prepared by ligating the full-length coding sequences of mouse Lamp1, mouse WASP, and human WAVE2 into a pMSCV retroviral expression vector upstream of GFP (*57*). The Lifeact-mApple fusion was prepared by PCR from an mApple-N1 template plasmid using oligos that encoded the Lifeact peptide N-terminal to mApple, followed by subcloning into pMSCV. shRNA constructs (shNT and shPTEN) (*8*) were subcloned into the miR30E vector (*58*) using the following primers: miRE-Xho-fw (5’- TGAACTCGAGAAGGTATAT TGCTGTTGACAGTGAGCG −3’) and miRE-EcoOligo-rev (5’-TCTCGAATTCT AGCCCCTTGAAGTCCGAGGCAGTAGGC −3’). gRNAs targeting WASP and WAVE2 were generated as previously described (*59, 60*) using the following oligos: NT gRNA: oligo-1 (5’- CACCG**GGATACCTGGGCCGACTTTC** −3’) and oligo-2 (5’- AAAC**GAAAGTCGGCCCAGGTATCC**C −3’). WASP gRNA: oligo-1 (5’- CACCG**CTGGACCATGGAACACTGCG** −3’) and oligo-2 (5’- AAAC **CGCAGTGTTCCATGGTCCAG**C −3’). WAVE2 gRNA: oligo-1 (5’- CACCG**AGCAAGGGAGTTTACTCGGG** −3’) and oligo-2 (AAAC **CCCGAGTAAACTCCCTTGCT**C −3’). Each gRNA was subsequently cloned into the LentiGuide-Puro vector and amplified by PCR using the following primers: LMP BamHI F2 (5’- TTTTTGGATCCTAGTAGGAGGCTTGGTAG −3’) and LMP EcoRI R2 (5’-TTTTTGAATTCTGTCTACTATTCTTTCCC −3’). The fragments were then digested using BamHI and EcoRI and ligated into miR30E previously digested with EcoRI and BglII.

### Cells and small molecule inhibitors

The animal protocols used for this study were approved by the Institutional Animal Care and Use Committee of Memorial Sloan-Kettering Cancer Center. Primary CTL blasts were prepared by pulsing irradiated C57BL/6 splenocytes with 100 nM OVA and then mixing them with T cells from OT1 αβTCR transgenic mice (Taconic) in RPMI medium containing 10% (vol/vol) FCS. Cells were supplemented with 30 IU/ml IL2 after 24 h and were split as needed in RPMI medium containing 10% (vol/vol) FCS and IL2. RMA-s cells were maintained in RPMI containing 10% (vol/vol) FCS while C57BL/6 murine cardiac microvascular endothelial cells (CellBiologics) were maintained in DMEM medium containing 10% (vol/vol) FCS. Inhibition of the Arp2/3 complex was achieved by preincubation of CTLs with CK666 (100-150 μM, Sigma-Aldrich) for 10 min at 37°C. The CTLs were then maintained in presence of CK666 for the duration of the experiment.

### Retroviral transduction

Phoenix E cells were transfected with expression vectors together with packaging plasmids using the calcium phosphate method. Ecotropic viral supernatants were collected after 48 h at 37°C and added to 1.5 × 10^6^ OT1 blasts 2 days after primary peptide stimulation. Mixtures were centrifuged at 1400 × g in the presence of polybrene (4 μg/ml) at 35 ^o^C. T cells were then split 1:3 in RPMI medium containing 10% (vol/vol) FCS and IL2 and allowed to grow for an additional 4-6 days.

### Micropillar preparation

PDMS (Sylgard 184; Dow Corning) micropillar arrays were prepared as previously described (*17*). Two types of pillars were used for this study: 1) 1 μm diameter, 5 μm in height and spaced hexagonally with a 2 μm center-to-center distance (thick pillars used for imaging protrusions), and 2) 0.7 μm diameter, 6 μm in height and spaced hexagonally with a 2 μm center-to-center distance (thin pillars used to measure synaptic forces). Micropillars were cast on chambered coverglass (Lab-Tek), washed with ethanol and Phosphate Buffered Saline (PBS), and stained with 20 μg/ml fluorescently-labeled streptavidin (Alexa647 or Alexa568, ThermoFisher Scientific) for 2 h at room temperature. After additional washing in PBS, the arrays were incubated with biotinylated H2-K^b^-OVA and ICAM1 (10 μg/ml each) O/N at 4 ^o^C (*8*). The pillars were then washed into RPMI containing 5% (vol/vol) FCS and lacking phenol red for imaging. Pillars prepared in this manner appear uniformly fluorescent in imaging experiments, indicating that the protein coating is homogenous.

### Live imaging on micropillars

For force measurement experiments, T cells were stained with Alexa488-labeled anti-CD45.2 F_ab_ (clone 104-2, MSKCC Monoclonal Antibody Core) and imaged on fluorescently labeled 0.7 μm diameter pillars. Videomicroscopy was performed using an inverted fluorescence microscope (Olympus IX-81) fitted with a 100 × objective lens (Olympus) and a mercury lamp for excitation. Images in the 488 nm (CTLs) and 568 nm (pillars) channels were collected every 15 s using Metamorph software. Protrusion formation was imaged on 1 μm diameter pillars stained using Alexa647-labeled streptavidin. Cells expressing fluorescent probes were added to the arrays and imaged using a confocal laser scanning microscope fitted with a 40 × objective lens and 488 nm, 560 nm, and 642 nm lasers (Leica SP5 or Zeiss LSM 880).

### Lattice light-sheet imaging

Lattice light-sheet microscopy was performed as described previously using 488 nm, 560 nm, and 642 nm lasers for illumination and a 25 × water immersion objective lens (*61*). Micropillar arrays were cast on 5 mm diameter coverslips, which were coated with stimulatory proteins as described above and then mounted for imaging. Videos (3-20 s time-lapse intervals) were recorded immediately after addition of fluorescently labeled CTLs using two cameras. 488 nm (30-60 mW laser power) and 560 nm (50 mW laser power) images were collected on one camera and 642 nm (50 mW laser power) images on a second camera. For CTL-target cell imaging, CTLs were added to coverslips bearing 90% confluent monolayers of iRFP670-labeled endothelial cells that had been incubated overnight in 2 μM OVA. Videos (15-20 s time-lapse intervals) were recorded immediately after addition of CTLs. 488 nm (50 mW laser power), 560 nm (200 mW laser power), and 642 nm (200 mW laser power) images were collected on one camera. Raw data were deskewed and deconvolved as described (*61*) using experimentally derived point spread functions. For two camera experiments, image alignment was performed in Matlab using reference images of fluorescent beads.

### Fixed imaging

OT1 cells were incubated on stimulatory micropillars for 20 min at 37°C, fixed by adding 4% paraformaldehyde for 5 min and washed with PBS. Samples were then blocked in PBS solution supplemented with 2% goat serum for 1 h at room temperature and incubated overnight at 4 °C using anti-β-tubulin (clone TUB 2.1; Sigma) or anti-LFA1 (clone M14/7; eBioscience). Actin was stained using Alexa 546-labeled phalloidin (ThermoFisher Scientific). After washing, samples were incubated with the appropriate secondary antibody for 2h at room temperature, washed and imaged using a Leica SP8 confocal laser scanning microscope fitted with a white light laser and a 40 × objective lens.

### Ca^2+^ imaging

CTLs were loaded with 5 μg/ml Fura2-AM (ThermoFisher Scientific), washed, and then imaged on stimulatory glass surfaces coated with H-2K^b^-OVA and ICAM-1 as previously described (*57*). Images were acquired using 340 nm and 380 nm excitation every 30 seconds for 30 min with a 20 × objective lens (Olympus).

### Image analysis

Imaging data were analyzed using SlideBook (3I), Imaris (Bitplane), Excel (Microsoft), Prism (GraphPad), and Matlab (MathWorks). Ca^2+^ responses were quantified by first normalizing the ratiometric Fura2 response of each individual cell to the last time point before the initial influx of Ca^2+^, and then by aligning and averaging all responses in the data set based on this initial time of influx. To quantify F-actin intensity in fixed images (fig. S3C), the sum intensity of Alexa546-labeled phalloidin in the region beneath the pillar tops was determined for each cell after intensity thresholding. For protrusion enrichment (Fig. 3C), total Lifeact-GFP intensity in the region beneath the pillar tops (F_A_) was divided by the total Lifeact-GFP intensity in a region of identical volume beginning from the first z-section above the pillar tops and extending upwards (F_B_) (fig. S4A). Force exertion against 0.7 μm diameter pillars was calculated after extracting pillar displacements from the imaging data and then converting these displacements into force vectors using custom Matlab scripts (*17, 18*). Radial distributions of pillar deflections (Fig. 6C) were generated by calculating the distances between strongly deflected pillars (> 0.6 μm deflection) and the IS COG at each time point. COG coordinates for the IS were generated from masks derived from the Alexa488 (CTL) channel (fig. S5B). To calculate the centralization factor (Fig. 1D, 5B, 5D), a mask encompassing the entire IS (M_C_) and a mask containing only the features of interest (e.g. lytic granules) (M_F_) were first generated using xy projection images. Then, the average distance between every pixel in M_C_ and the COG of M_C_ (D_C_) was determined and subsequently divided by the average distance between every pixel in M_F_ and the COG of M_C_ (D_F_) (fig. S2). Granule polarization (Fig. 1D) was quantified by determining the vertical distance between the centroid of a mask encompassing the lytic granules and the deepest CTL protrusion, using xz or yz projection images. To calculate target IS volume (Fig. 7B and C), a three dimensional mask was generated at a time point of interest by tracing the edges of the IS and then propagating the resulting shape downward to encompass the sample lying directly beneath the CTL. The volume of this region occupied by the endothelial cell (V_1_) was then divided by the volume of this same region occupied by the endothelial cell at time 0 (V_0_) (fig. S6D).

### Adhesion assay

Integrin-mediated adhesion by OT1 CTLs was measured in flat-bottomed 96-well plates bearing increasing densities of ICAM1. The plates were coated with 10 μg/ml streptavidin in PBS followed by increasing concentrations of biotinylated ICAM1 in the presence or absence of 1 μg/ml biotinylated H2-K^b^-OVA in Hepes buffered saline (10 mM Hepes pH 7.5, 150 mM NaCl) with 2% BSA. The total concentration of biotinylated protein during coating was kept at 5 μg/ml by the addition of nonspecific biotinylated MHC (H2-D^b^). CTLs were fluorescently labeled with cell trace violet (CTV), resuspended in adhesion buffer (PBS with 0.5% BSA, 2 mM MgCl_2_, 1 mM CaCl_2_), and added to ICAM1-bearing wells in triplicate. After a 20 min incubation at 37 °C, wells were washed with warmed adhesion buffer as described (*62*), and the bound cells quantified by fluorimetry.

### Killing assay, degranulation and conjugate formation

RMA-s target cells were labeled with carboxyfluorescein succinimidyl ester (CFSE) or the membrane dye PKH26, loaded with different concentrations of OVA and mixed in a 96-well V-bottomed plate with CTV stained OT-1 cells. To assess killing, cells were mixed at a 1:3 E:T ratio and incubated for 4 h at 37 °C. Specific lysis of CFSE^+^ target cells was determined by flow cytometry as previously described (*63*). For degranulation assays, the E:T ratio was 1:1 and cells were incubated at 37 °C for 90 min in the presence of eFluor 660-labeled Lamp1 Monoclonal Antibody (clone eBio1D4B, eBioscience). Lamp1 staining was then assessed by flow cytometry. To measure conjugate formation, CTLs and targets were mixed 1:1, lightly centrifuged (100 × g) to encourage cell contact, and incubated for 20 min at 37 °C. Then, cells were then resuspended in the presence of 2% PFA, washed in FACS buffer (PBS + 4% FCS), and analyzed by flow cytometry. Conjugate formation was quantified as (CFSE^+^ CTV^+^)/(CTV^+^).

### Immunoblot

0.2-1 × 10^6^ CTLs were lysed using cold cell lysis buffer containing 50 mM TrisHCl, 0.15 M NaCl, 1 mM EDTA, 1% NP-40 and 0.25% sodium deoxycholate. Suppression of PTEN, WASP and WAVE2 was confirmed using the following antibodies: anti-PTEN monoclonal antibody (clone D4.3; Cell Signaling Technology), anti-WASP monoclonal antibody (clone B-9; Santa Cruz) and anti-WAVE2 monoclonal antibody (clone D2C8; Cell Signaling Technology). Actin served as a loading control (clone AC-15, Sigma). For signaling assays, serum and IL2 starved OT1 CTLs were incubated with streptavidin polystyrene beads (Spherotech) coated with H2-K^b^-OVA and ICAM1 at a 1:1 ratio for various times at 37 °C and immediately lysed in 2 × cold lysis buffer containing phosphatase inhibitors (1 mM NaF and 0.1 mM Na_3_VO_4_) and protease inhibitors (cOmplete mini cocktail, EDTA-free, Roche). Activation of PI3K, MAP kinase and NF-κB signaling was assessed by immunoblot for pAkt (Phospho-Akt (Ser473) Ab; Cell Signaling Technology), pErk1/2 (Phospho-Thr202/ Tyr204; clone D13.14.4E; Cell Signaling Technology), and IκB (Cell Signaling Technology).

### Statistical analysis

Figures show representative experiments. Analysis was carried out using either representative experiments or pooled data as indicated (N refers to the number of cells analyzed). Statistical analyses (unpaired or paired T tests) were carried out using GraphPad Prism or Microsoft Excel. All error bars denote SEM.

## ACKNOWLEDGMENTS

We thank A. Kepecs, C. Firl, L. Foulon, and T. Lipsky for technical support; T. L. Chew, J. Aaron, S. Khuon, and E. Wait for assistance with lattice light-sheet imaging; Y. Romin, S. Fujisawa, E. Feng, and the MSKCC Molecular Cytology Core Facility for assistance with confocal imaging; S. Lowe for the miR30E vector; the MSKCC Monoclonal Antibody Core Facility for fluorescently conjugated F_ab_ fragments; S. Rudensky for critical assessment of the manuscript; and members of the M. H. and L. C. K. labs for advice. This work was supported in part by the NIH (R01-AI087644 to M. H., R01-AI110593 to L. C. K., P30-CA008748 to MSKCC), the NSF (CMMI-1562905 to M. H. and L. C. K.), the Leukemia and Lymphoma Society (M. H.), the Cancer Research Institute (F. T.), and HHMI (J. M. H.).

## SUPPLEMENTARY MATERIALS

**Figure S1.**
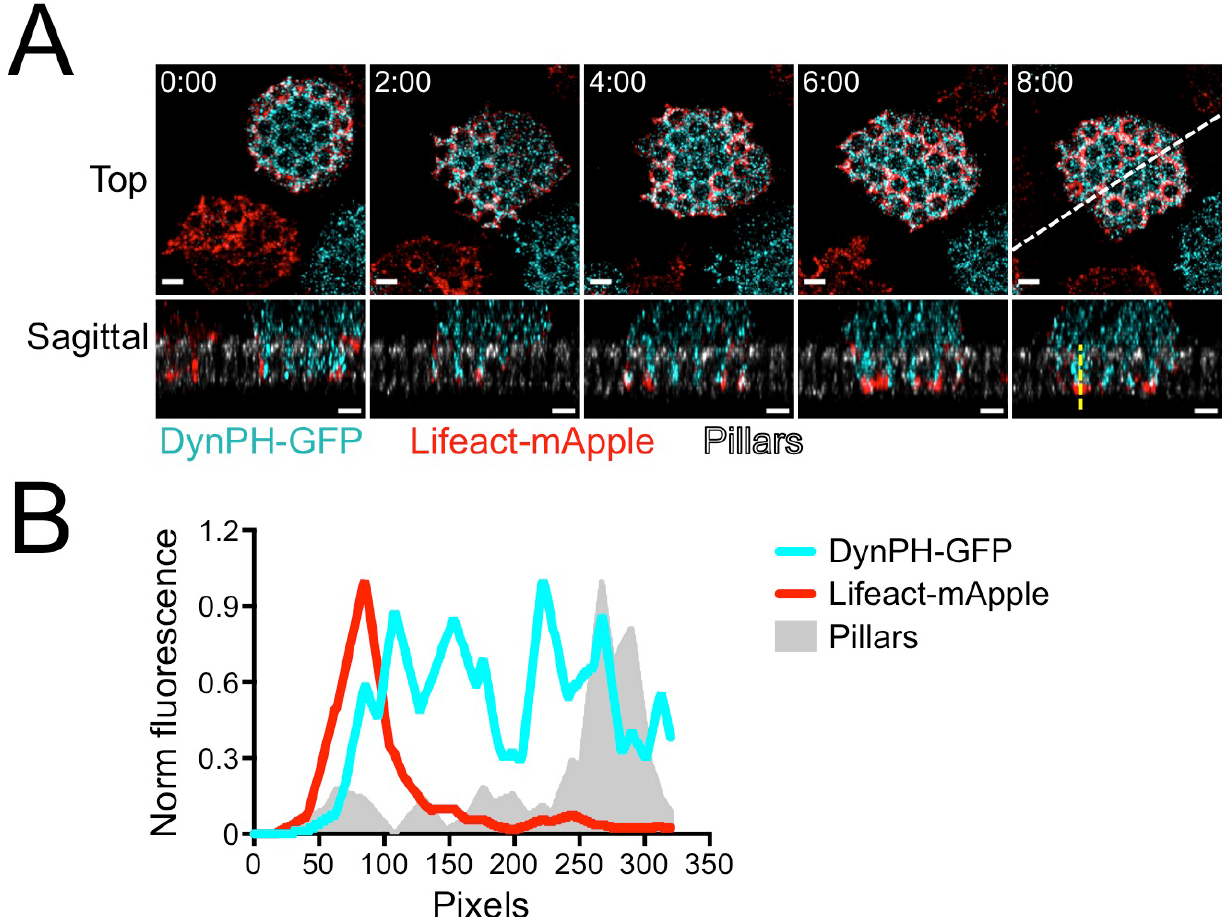
F-actin accumulates at the leading edges of synaptic protrusions. OT1 CTLs expressing Lifeact-mApple and DynPH-GFP (a membrane marker) were imaged by confocal microscopy on fluorescent micropillars bearing H2-K^b^-OVA and ICAM1. (A) Time-lapse montage of a representative CTL, with z-projection images (top views) shown above and corresponding sagittal views below. The white dotted line denotes the slicing plane used for the sagittal images. Pillars (gray) appear in the sagittal images only. Time in M:SS is indicated in the upper left corner of each top view. Scale bars = 2 μm. (B) Linescan (derived from the dotted yellow line in A) showing normalized fluorescence intensity of DynPH-GFP, Lifeact-mApple, and the pillars.

**Figure S2.**
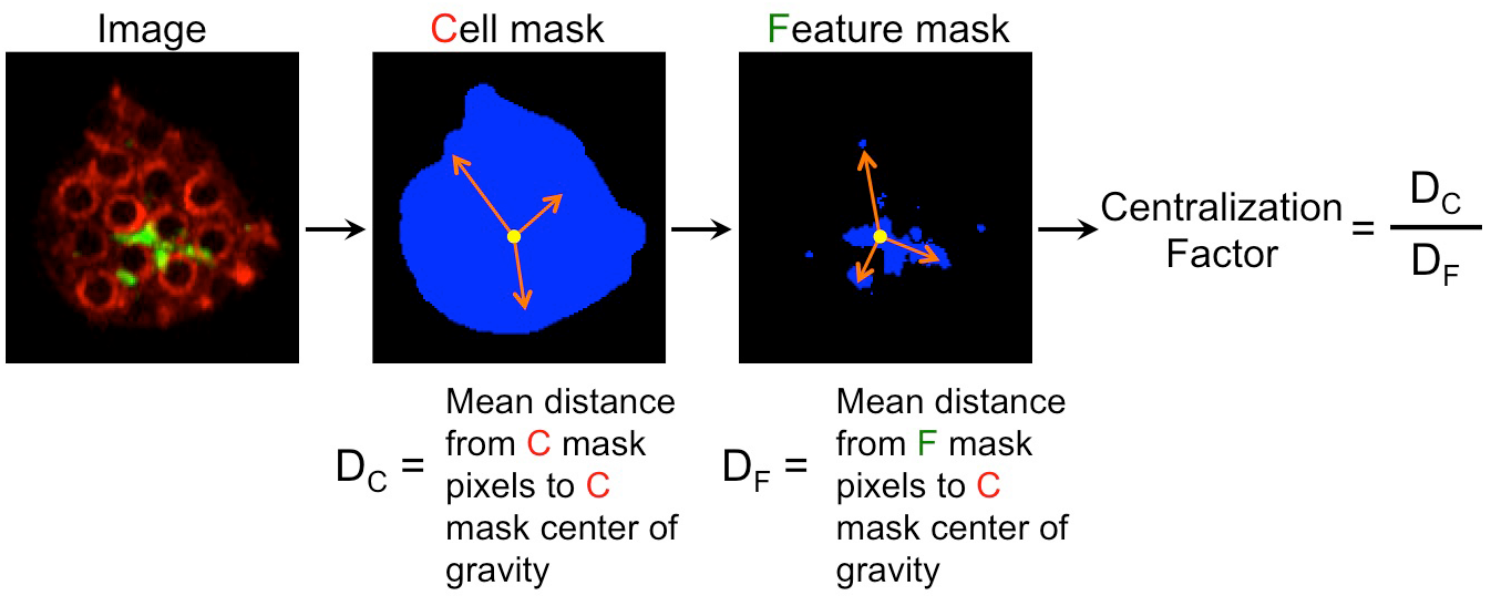
Centralization factor analysis. The centralization factor compares the mean distance between a given feature within the IS (e.g. lytic granules) and the IS center of gravity (D_F_) to the mean distance between all positions within the IS and the IS center of gravity (D_C_).

**Figure S3.**
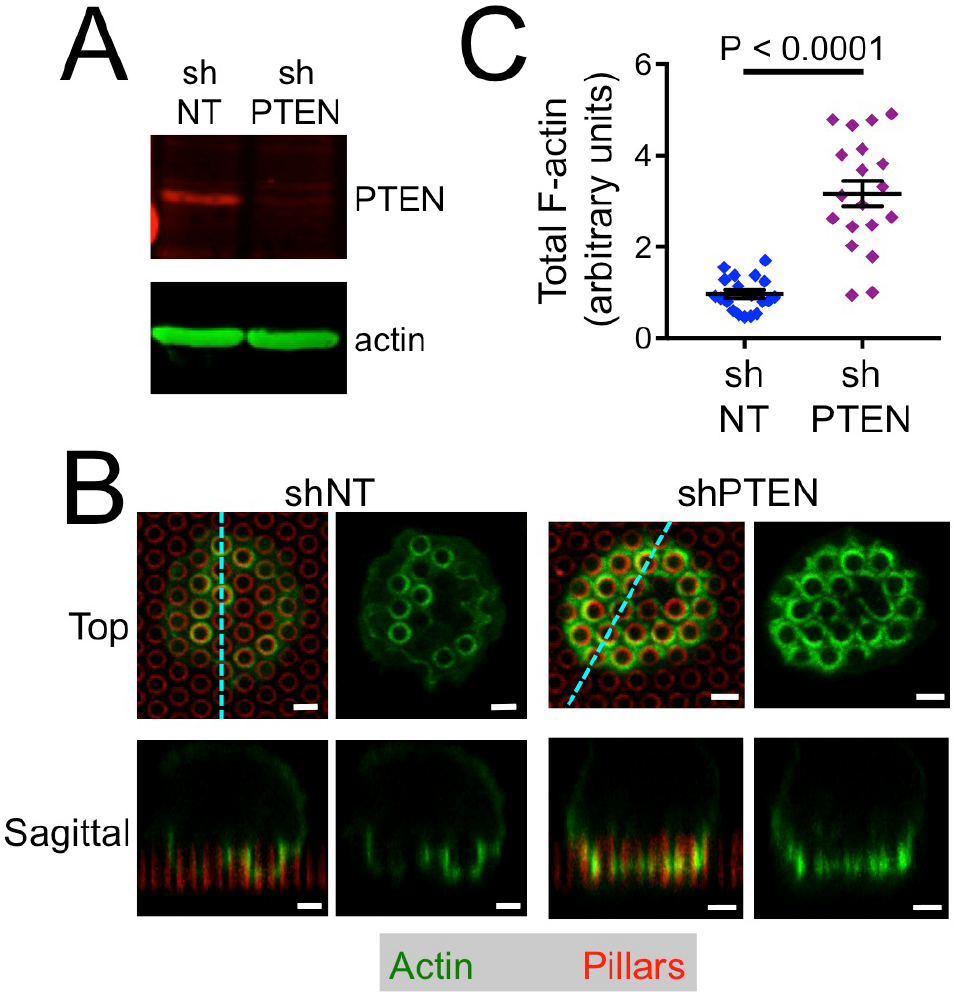
PTEN depletion enhances F-actin accumulation in protrusions. OT1 CTLs expressing shRNA against PTEN (shPTEN) or a control nontargeting shRNA (shNT) were applied to fluorescent micropillars bearing H2-K^b^-OVA and ICAM1, fixed, and stained with phalloidin (to visualize F-actin). (A) Immunoblot analysis of PTEN expression in shNT and shPTEN CTLs. Actin served as a loading control. (B) Representative images of shNT and shPTEN CTLs, with z-projection images (top views) shown above and corresponding sagittal views below. Cyan dotted lines denote the slicing planes used for the sagittal images. Scale bars = 2 μm. (C) Quantification of sum F-actin intensity on micropillar arrays. Error bars denote SEM. N = 19 for each cell type. P calculated from two-tailed Student’s T-test.

**Figure S4.**
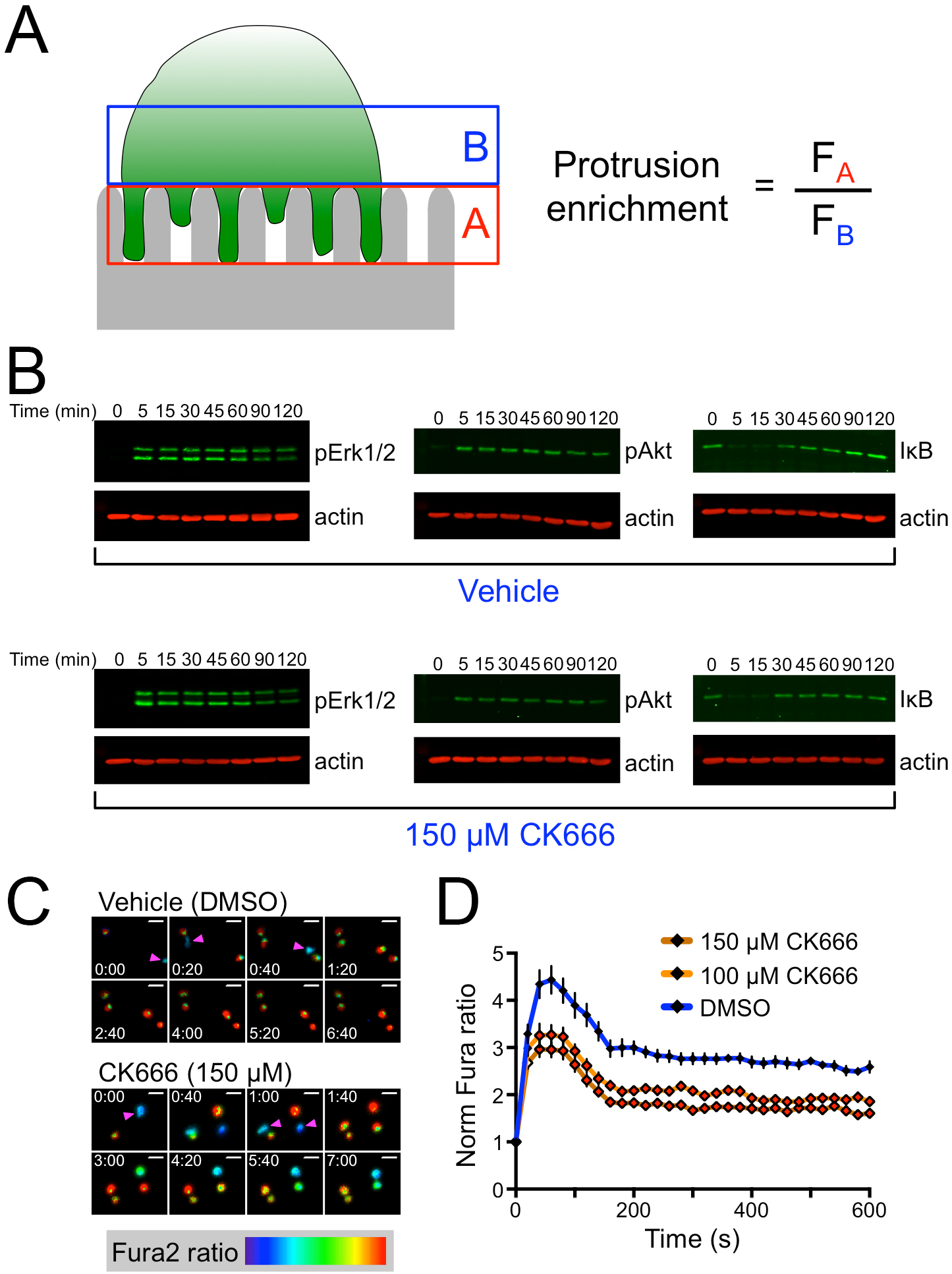
Effects of CK666 on T cell activation. (A) Diagram schematizing the quantification of synaptic protrusions shown in Fig. 3C. (B) OT1 CTLs were stimulated with beads coated with H2-K^b^-OVA and ICAM1 in the presence of 150 μM CK666 or DMSO vehicle for the indicated times and then lysed. pErk1/2, pAKT, and IκB levels were assessed by immunoblot using actin as a loading control. (C-D) OT1 CTLs were loaded with Fura2-AM Ca^2+^ dye, incubated with the indicated concentrations of CK666 or DMSO vehicle, and then imaged on stimulatory glass surfaces coated with H2-K^b^-OVA and ICAM1. (C) Time-lapse montages of representative Ca^2+^ responses, with Fura2 ratio displayed in pseudocolor (warmer colors denote higher intracellular Ca^2+^ concentration). Magenta arrowheads highlight CTLs at the time point just before the onset of Ca^2+^ flux. Time in M:SS is indicated in each image. Scale bars = 20 μm. (D) Mean normalized Fura2 ratio was graphed against time (see Materials and Methods). N = 30 for each cell type, with error bars denoting SEM. Filled white diamonds and filled red diamonds indicate P < 0.05 and P < 0.01, respectively, calculated by two-tailed Student’s T-test comparing each CK666 condition against DMSO control.

**Figure S5.**
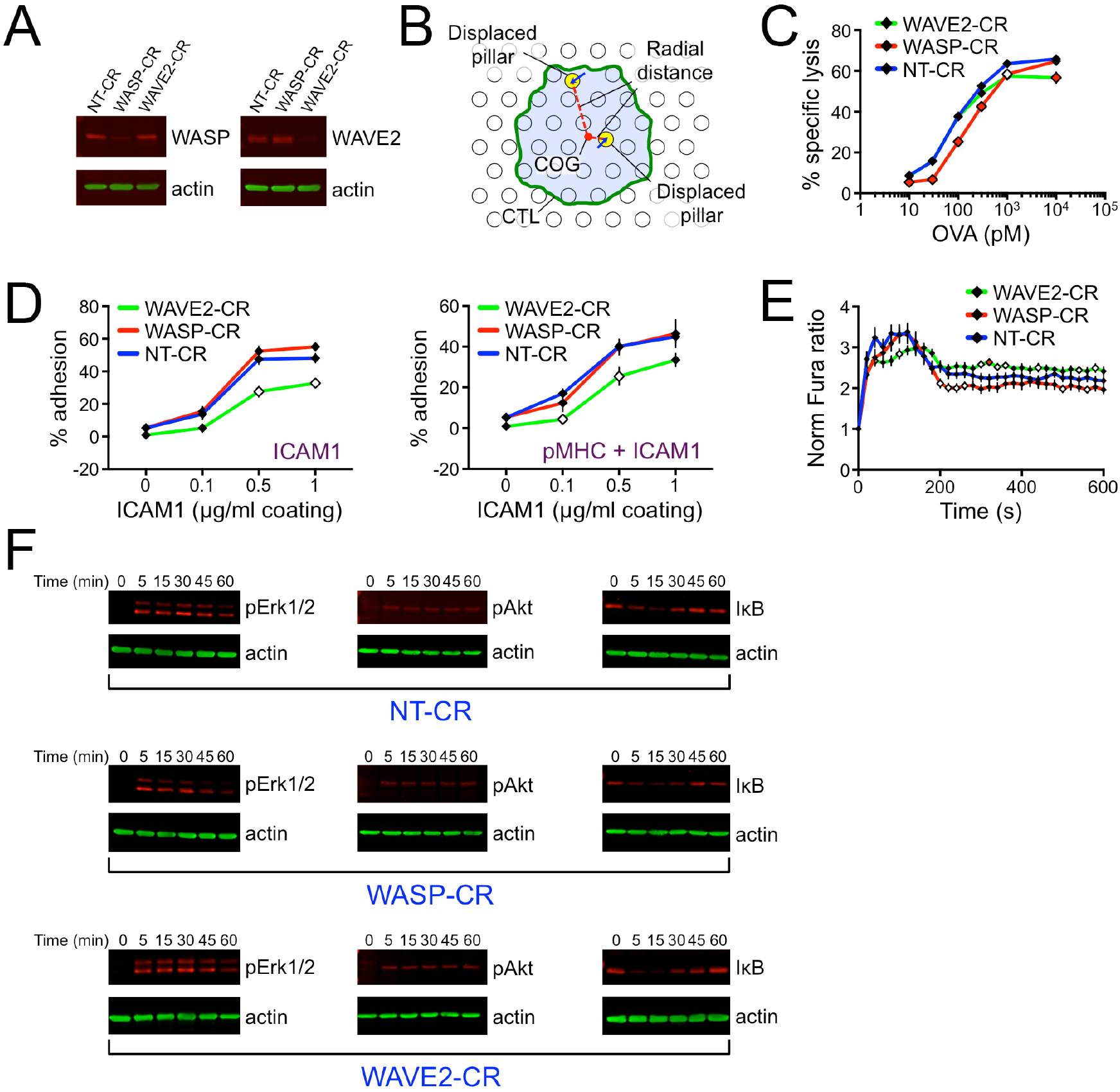
WASP and WAVE2 depletion induce distinct functional phenotypes. (A) Immunoblot analysis of WASP and WAVE2 expression in NT-CR, WASP-CR, and WAVE2-CR CTLs. Actin served as a loading control. (B) Diagram schematizing the radial distance histogram analysis of pillar deflections shown in Fig. 6C. (C) RMA-s target cells were loaded with increasing concentrations of OVA and then mixed with NT-CR, WASP-CR, or WAVE2-CR CTLs. Specific lysis of RMA-s cells is shown, with error bars denoting SEM. This experiment highlights the cytotoxicity defect occasionally observed in WAVE2-CR CTLs at high OVA concentrations. (D) Adhesion of fluorescently labeled NT-CR, WASP-CR, and WAVE2-CR OT1 CTLs to wells coated with the indicated concentrations of ICAM1 in the absence (left) or presence (right) of H2-K^b^-OVA (pMHC). Error bars denote SEM. (E) NT-CR, WASP-CR, and WAVE2-CR OT1 CTLs were loaded with Fura2-AM Ca^2+^ dye and then imaged on stimulatory glass surfaces coated with H2-K^b^-OVA and ICAM1. Ca^2+^ signaling was analyzed by quantifying mean normalized Fura2 ratio over time (see Materials and Methods). N ≥ 29 for each cell type, with error bars denote SEM. In C, D, and E, filled white diamonds and filled red diamonds indicate P < 0.05 and P < 0.01, respectively, calculated by two-tailed Student’s T-test comparing WASP-CR and WAVE2-CR to NT-CR. (F) NT-CR, WASP-CR, and WAVE2-CR OT1 CTLs were stimulated with beads coated with H2-K^b^-OVA and ICAM1 for the indicated times and then lysed. pErk1/2, pAKT, and IκB levels were assessed by immunoblot using actin as a loading control.

**Figure S6.**
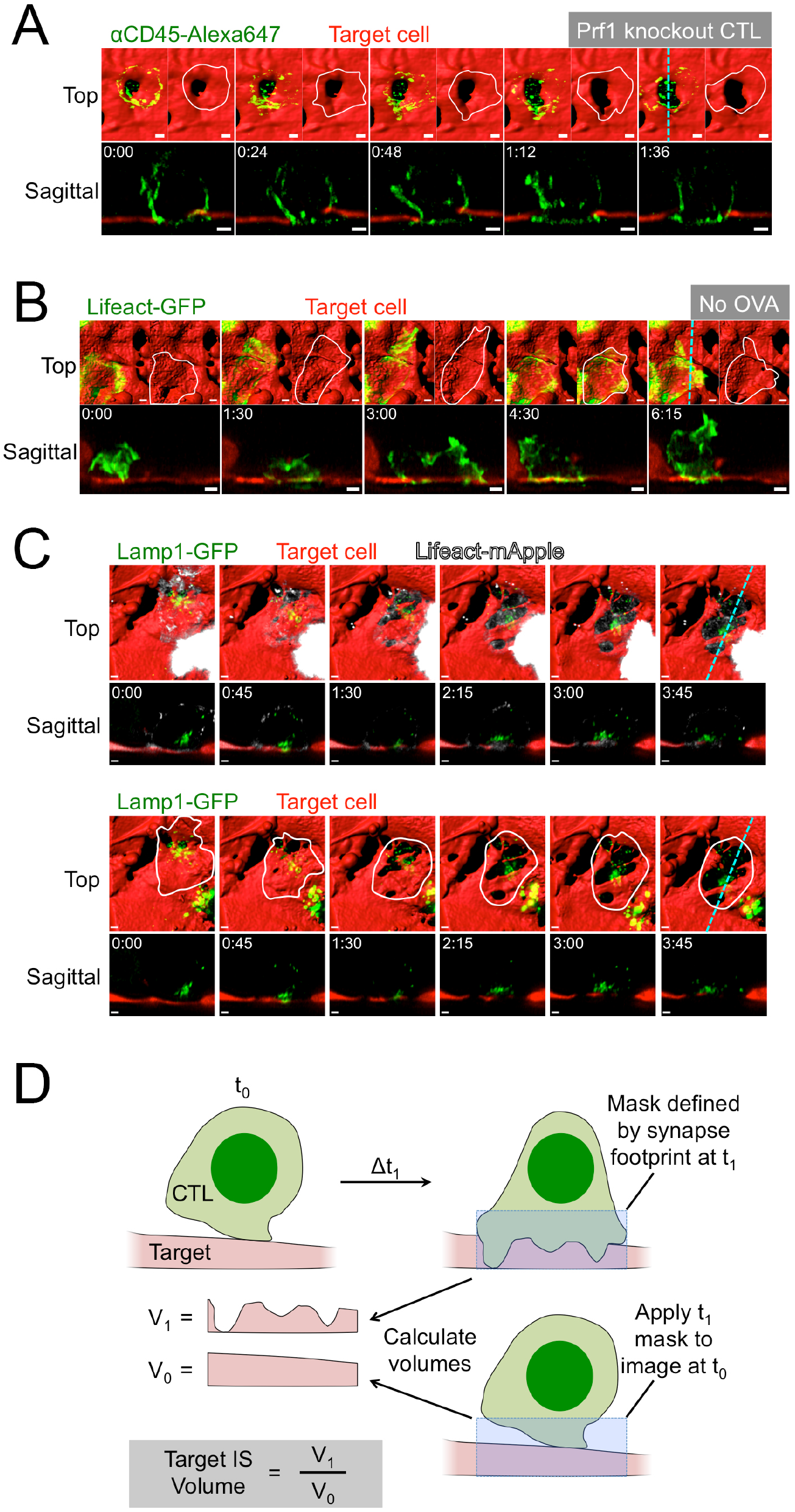
CTLs physically distort the target cell surface. (A-C) OT1 CTLs were applied to confluent cultures of OVA-loaded endothelial target cells expressing mApple (A) or iRFP670 (B and C) and then imaged using lattice light-sheet microscopy. All time-lapse montages show “vertically” oriented synapses, with z-projection images (top views) shown above and corresponding sagittal views below. Target cells are visualized by surface representation in all z-projection images. Cyan dotted lines denote the slicing planes used for the sagittal images. Time in M:SS is indicated in the upper left corner of each sagittal image. Scale bars = 2 μm. (A) A representative *Prf1*^−/−^ CTL, visualized using fluorescent F_ab_ fragments against CD45, engaging a target cell beneath it. (B) A representative CTL expressing Lifeact-GFP engaging a target cell in the absence of OVA. In A and B, two z-projections are shown for each time point; the fluorescent signal from the CTL appears on the left and is replaced with an outline of the CTL of interest on the right. (C) A representative CTL expressing Lifeact-mApple and Lamp1-GFP, engaging a target cell beneath it. The montage above includes the Lifeact-mApple signal, whereas the montage below contains an outline of the CTL of interest. (D) The displacement index compares the endothelial cell volume in the region beneath a CTL at a given time (V_1_) with the volume occupied by the endothelial cell in that same region at time 0 (V_0_).

## SUPPLEMENTAL VIDEO LEGENDS

**Video S1. CTLs form dynamic F-actin rich protrusions on stimulatory micropillars.** OT1 CTLs expressing Lifeact-GFP were imaged using lattice light-sheet microscopy on thick PDMS micropillar arrays coated with H2-K^b^-OVA and ICAM1. A 60 × time-lapse video is shown of a representative CTL landing on the array. A z-projection (top view) of the CTL appears on the left and a sagittal slice on the right. Pillars are not visible but their shape can be inferred from the pattern of the CTL response. Time in H:MM:SS is shown at the bottom left. Scale bar = 2 μm.

**Video S2. CTL degranulation occurs at the base of synaptic protrusions.** OT1 CTLs expressing Lifeact-mRuby2 (red) and pHluorin-Lamp1 (cyan) were imaged using lattice light-sheet microscopy on thick PDMS micropillar arrays coated with H2-K^b^-OVA and ICAM1. A 30 × time-lapse video is shown of a representative degranulating CTL. A z-projection (top view) of the CTL appears on the left and a sagittal slice on the right. Pillars (gray) are shown in the sagittal view only. The degranulation event appears in the third frame of the video. Time in M:SS is shown at the bottom left. Scale bars = 2 μm.

**Video S3. Vehicle-treated CTLs form F-actin rich protrusions on stimulatory micropillars.** OT1 CTLs expressing Lifeact-GFP (green) were treated with DMSO (vehicle control) and imaged using confocal microscopy on thick PDMS micropillar arrays coated with H2-K^b^-OVA and ICAM1. A 125 × time-lapse video is shown of a representative CTL landing on the array. A z-projection (top view) of the CTL appears on the left and a sagittal slice on the right. Pillars are shown in red. Time in M:SS is shown at the bottom left. Scale bar = 2 μm.

**Video S4. CK666-treated CTLs form F-actin rich protrusions on stimulatory micropillars.** OT1 CTLs expressing Lifeact-GFP (green) were treated with 150 μM CK666 and imaged using confocal microscopy on thick PDMS micropillar arrays coated with H2-K^b^-OVA and ICAM1. A 95 × time-lapse video is shown of a representative CTL landing on the array. A z-projection (top view) of the CTL appears on the left and a sagittal slice on the right. Pillars are shown in red. Time in H:MM:SS is shown at the bottom left. Scale bar = 2 μm.

**Video S5. Force exertion by vehicle-treated CTLs.** OT1 CTLs were stained with an Alexa488-labeled anti-CD45 F_ab_ fragment, treated with DMSO (vehicle control), and imaged on thin PDMS micropillar arrays coated with H2-K^b^-OVA and ICAM1. A 75 × time-lapse video is shown of a representative CTL in the plane of the pillar tops. Pillars appear on the left, the CTL in the center, and the overlay image on the right. Time in M:SS is shown at the bottom right.

**Video S6. Force exertion by CK666-treated CTLs.** OT1 CTLs were stained with an Alexa488-labeled anti-CD45 F_ab_ fragment, treated with 150 μM CK666, and imaged on thin PDMS micropillar arrays coated with H2-K^b^-OVA and ICAM1. A 75 × time-lapse video is shown of a representative CTL in the plane of the pillar tops. Pillars appear on the left, the CTL in the center, and the overlay image on the right. Time in M:SS is shown at the bottom right.

**Video S7. WAVE2-GFP accumulates in peripheral synaptic protrusions.** OT1 CTLs expressing WAVE2-GFP (green) were imaged using confocal microscopy on thick PDMS micropillar arrays coated with H2-K^b^-OVA and ICAM1. A 150 × time-lapse video is shown of a representative CTL landing on the array. A z-projection (top view) of the CTL appears on the left and a sagittal slice on the right. Pillars are shown in red. Time in H:MM:SS is shown at the bottom left. Scale bars = 2 μm.

**Video S8. WASP-GFP accumulates in central synaptic protrusions.** OT1 CTLs expressing WASP-GFP (green) were imaged using confocal microscopy on thick PDMS micropillar arrays coated with H2-K^b^-OVA and ICAM1. A 150 × time-lapse video is shown of a representative CTL landing on the array. A z-projection (top view) of the CTL appears on the left and a sagittal slice on the right. Pillars are shown in red. Time in H:MM:SS is shown at the bottom left. Scale bars = 2 μm.

**Video S9. Protrusive behavior of NT-CR CTLs on stimulatory micropillar arrays.** NT-CR OT1 CTLs expressing Lifeact-GFP (green) were imaged using confocal microscopy on thick PDMS micropillar arrays coated with H2-K^b^-OVA and ICAM1. A 75 × time-lapse video is shown of a representative CTL landing on the array. A z-projection (top view) of the CTL appears on the left and a sagittal slice on the right. Pillars are shown in red. Time in M:SS is shown at the bottom. Scale bars = 2 μm.

**Video S10. Protrusive behavior of WASP-CR CTLs on stimulatory micropillar arrays.** WASP-CR OT1 CTLs expressing Lifeact-GFP (green) were imaged using confocal microscopy on thick PDMS micropillar arrays coated with H2-K^b^-OVA and ICAM1. A 75 × time-lapse video is shown of a representative CTL landing on the array. A z-projection (top view) of the CTL appears on the left and a sagittal slice on the right. Pillars are shown in red. Time in M:SS is shown at the bottom. Scale bars = 2 μm.

**Video S11. Protrusive behavior of WAVE2-CR CTLs on stimulatory micropillar arrays.** WAVE2-CR OT1 CTLs expressing Lifeact-GFP (green) were imaged using confocal microscopy on thick PDMS micropillar arrays coated with H2-K^b^-OVA and ICAM1. A 75 × time-lapse video is shown of a representative CTL landing on the array. A z-projection (top view) of the CTL appears on the left and a sagittal slice on the right. Pillars are shown in red. Time in M:SS is shown at the bottom. Scale bars = 2 μm.

**Video S12. Force exertion by NT-CR CTLs.** NT-CR OT1 CTLs were stained with an Alexa488-labeled anti-CD45 F_ab_ fragment and imaged on thin PDMS micropillar arrays coated with H2-K^b^-OVA and ICAM1. A 75 × time-lapse video is shown of a representative CTL in the plane of the pillar tops. Pillars appear on the left, the CTL in the center, and the overlay image on the right. Time in M:SS is shown at the bottom right.

**Video S13. Force exertion by WASP-CR CTLs.** WASP-CR OT1 CTLs were stained with an Alexa488-labeled anti-CD45 F_ab_ fragment and imaged on thin PDMS micropillar arrays coated with H2-K^b^-OVA and ICAM1. A 75 × time-lapse video is shown of a representative CTL in the plane of the pillar tops. Pillars appear on the left, the CTL in the center, and the overlay image on the right. Time in M:SS is shown at the bottom right.

**Video S14. Force exertion by WAVE2-CR CTLs.** WAVE2-CR OT1 CTLs were stained with an Alexa488-labeled anti-CD45 F_ab_ fragment and imaged on thin PDMS micropillar arrays coated with H2-K^b^-OVA and ICAM1. A 75 × time-lapse video is shown of a representative CTL in the plane of the pillar tops. Pillars appear on the left, the CTL in the center, and the overlay image on the right. Time in M:SS is shown at the bottom right.

**Video S15. Target cell distortion by NT-CR CTLs.** NT-CR CTLs expressing Lifeact-GFP (green) were imaged using lattice light-sheet microscopy on endothelial target cells that had been loaded with OVA peptide. A 75 × time-lapse video is shown of a representative CTL attacking a target cell (red) beneath it. A z-projection (top view) of the CTL and target cell is shown above and a sagittal slice of the two cells below. In z-projection images, the target cell has been rendered in surface representation. In the images to the right, the CTL has been removed to highlight target cell distortion. Time in M:SS is shown at the top. Scale bars = 2 μm.

**Video S16. Target cell distortion by WASP-CR CTLs.** WASP-CR CTLs expressing Lifeact-GFP (green) were imaged using lattice light-sheet microscopy on endothelial target cells that had been loaded with OVA peptide. A 75 × time-lapse video is shown of a representative CTL attacking a target cell (red) beneath it. A z-projection (top view) of the CTL and target cell is shown above and a sagittal slice of the two cells below. In z-projection images, the target cell has been rendered in surface representation. In the images to the right, the CTL has been removed to highlight target cell distortion. Time in M:SS is shown at the top. Scale bars = 2 μm.

**Video S17. Target cell distortion by WAVE2-CR CTLs.** WAVE2-CR CTLs expressing Lifeact-GFP (green) were imaged using lattice light-sheet microscopy on endothelial target cells that had been loaded with OVA peptide. A 75 × time-lapse video is shown of a representative CTL attacking a target cell (red) beneath it. A z-projection (top view) of the CTL and target cell is shown above and a sagittal slice of the two cells below. In z-projection images, the target cell has been rendered in surface representation. In the images to the right, the CTL has been removed to highlight target cell distortion. Time in M:SS is shown at the top. Scale bars = 2 μm.

## REFERENCES AND NOTES

1. F. D. Batista, M. L. Dustin, Cell:cell interactions in the immune system. Immunol Rev 251, 7–12 (2013).

2. M. L. Dustin, E. O. Long, Cytotoxic immunological synapses. Immunol Rev 235, 24–34 (2010).

3. J. C. Stinchcombe, G. M. Griffiths, Secretory mechanisms in cell-mediated cytotoxicity. Annual review of cell and developmental biology 23, 495–517 (2007).

4. J. Thiery, J. Lieberman, Perforin: a key pore-forming protein for immune control of viruses and cancer. Subcell Biochem 80, 197–220 (2014).

5. L. Martinez-Lostao, A. Anel, J. Pardo, How Do Cytotoxic Lymphocytes Kill Cancer Cells? Clin Cancer Res 21, 5047–5056 (2015).

6. X. Liu, M. Huse, in Cell Polarity 1: BIological Role and Basic Mechanisms, K. Ebnet, Ed. (Springer, Switzerland, 2015), pp. 247–275.

7. S. C. Bunnell, V. Kapoor, R. P. Trible, W. Zhang, L. E. Samelson, Dynamic actin polymerization drives T cell receptor-induced spreading: a role for the signal transduction adaptor LAT. Immunity 14, 315–329 (2001).

8. A. Le Floc’h, Y. Tanaka, N. S. Bantilan, G. Voisinne, G. Altan-Bonnet, Y. Fukui, M. Huse, Annular PIP3 accumulation controls actin architecture and modulates cytotoxicity at the immunological synapse. J Exp Med 210, 2721–2737 (2013).

9. Y. Kaizuka, A. D. Douglass, R. Varma, M. L. Dustin, R. D. Vale, Mechanisms for segregating T cell receptor and adhesion molecules during immunological synapse formation in Jurkat T cells. Proc Natl Acad Sci U S A 104, 20296–20301 (2007).

10. J. Yi, X. S. Wu, T. Crites, J. A. Hammer, 3rd, Actin retrograde flow and actomyosin II arc contraction drive receptor cluster dynamics at the immunological synapse in Jurkat T cells. Mol Biol Cell 23, 834–852 (2012).

11. J. C. Stinchcombe, E. Majorovits, G. Bossi, S. Fuller, G. M. Griffiths, Centrosome polarization delivers secretory granules to the immunological synapse. Nature 443, 462–465 (2006).

12. A. T. Ritter, Y. Asano, J. C. Stinchcombe, N. M. Dieckmann, B. C. Chen, C. Gawden-Bone, S. van Engelenburg, W. Legant, L. Gao, M. W. Davidson, E. Betzig, J. Lippincott-Schwartz, G. M. Griffiths, Actin depletion initiates events leading to granule secretion at the immunological synapse. Immunity 42, 864–876 (2015).

13. P. T. Sage, L. M. Varghese, R. Martinelli, T. E. Sciuto, M. Kamei, A. M. Dvorak, T. A. Springer, A. H. Sharpe, C. V. Carman, Antigen recognition is facilitated by invadosome-like protrusions formed by memory/effector T cells. J Immunol 188, 3686–3699 (2012).

14. E. Cai, K. Marchuk, P. Beemiller, C. Beppler, M. G. Rubashkin, V. M. Weaver, A. Gerard, T. L. Liu, B. C. Chen, E. Betzig, F. Bartumeus, M. F. Krummel, Visualizing dynamic microvillar search and stabilization during ligand detection by T cells. Science 356, (2017).

15. K. Nguyen, N. R. Sylvain, S. C. Bunnell, T cell costimulation via the integrin VLA-4 inhibits the actin-dependent centralization of signaling microclusters containing the adaptor SLP-76. Immunity 28, 810–821 (2008).

16. J. Husson, K. Chemin, A. Bohineust, C. Hivroz, N. Henry, Force generation upon T cell receptor engagement. PloS one 6, e19680 (2011).

17. K. T. Bashour, A. Gondarenko, H. Chen, K. Shen, X. Liu, M. Huse, J. C. Hone, L. Kam, CD28 and CD3 have complementary roles in T-cell traction forces. Proc Natl Acad Sci U S A 111, 2241–2246 (2014).

18. R. Basu, B. M. Whitlock, J. Husson, A. Le Floc’h, W. Jin, A. Oyler-Yaniv, F. Dotiwala, G. Giannone, C. Hivroz, N. Biais, J. Lieberman, L. C. Kam, M. Huse, Cytotoxic T Cells Use Mechanical Force to Potentiate Target Cell Killing. Cell 165, 100–110 (2016).

19. A. M. Beal, N. Anikeeva, R. Varma, T. O. Cameron, G. Vasiliver-Shamis, P. J. Norris, M. L. Dustin, Y. Sykulev, Kinetics of early T cell receptor signaling regulate the pathway of lytic granule delivery to the secretory domain. Immunity 31, 632–642 (2009).

20. J. C. Stinchcombe, G. Bossi, S. Booth, G. M. Griffiths, The immunological synapse of CTL contains a secretory domain and membrane bridges. Immunity 15, 751–761 (2001).

21. A. Kupfer, G. Dennert, Reorientation of the microtubule-organizing center and the Golgi apparatus in cloned cytotoxic lymphocytes triggered by binding to lysable target cells. J Immunol 133, 2762–2766 (1984).

22. G. D. Rak, E. M. Mace, P. P. Banerjee, T. Svitkina, J. S. Orange, Natural killer cell lytic granule secretion occurs through a pervasive actin network at the immune synapse. PLoS Biol 9, e1001151 (2011).

23. E. D. Goley, M. D. Welch, The ARP2/3 complex: an actin nucleator comes of age. Nat Rev Mol Cell Biol 7, 713–726 (2006).

24. T. S. Gomez, K. Kumar, R. B. Medeiros, Y. Shimizu, P. J. Leibson, D. D. Billadeau, Formins regulate the actin-related protein 2/3 complex-independent polarization of the centrosome to the immunological synapse. Immunity 26, 177–190 (2007).

25. S. Kumari, D. Depoil, R. Martinelli, E. Judokusumo, G. Carmona, F. B. Gertler, L. Kam, C. V. Carman, J. K. Burkhardt, D. J. Irvine, M. L. Dustin, Actin foci facilitate activation of the phospholipase C-gamma in primary T lymphocytes via the WASP pathway. eLife 4, (2015).

26. A. Le Floc’h, M. Huse, Molecular mechanisms and functional implications of polarized actin remodeling at the T cell immunological synapse. Cellular and molecular life sciences: CMLS 72, 537–556 (2015).

27. B. J. Nolen, N. Tomasevic, A. Russell, D. W. Pierce, Z. Jia, C. D. McCormick, J. Hartman, R. Sakowicz, T. D. Pollard, Characterization of two classes of small molecule inhibitors of Arp2/3 complex. Nature 460, 1031–1034 (2009).

28. S. Murugesan, J. Hong, J. Yi, D. Li, J. R. Beach, L. Shao, J. Meinhardt, G. Madison, X. Wu, E. Betzig, J. A. Hammer, Formin-generated actomyosin arcs propel T cell receptor microcluster movement at the immune synapse. J Cell Biol 215, 383–399 (2016).

29. H. L. Ostergaard, K. P. Kane, M. F. Mescher, W. R. Clark, Cytotoxic T lymphocyte mediated lysis without release of serine esterase. Nature 330, 71–72 (1987).

30. J. C. Nolz, T. S. Gomez, P. Zhu, S. Li, R. B. Medeiros, Y. Shimizu, J. K. Burkhardt, B. D. Freedman, D. D. Billadeau, The WAVE2 complex regulates actin cytoskeletal reorganization and CRAC-mediated calcium entry during T cell activation. Curr Biol 16, 24–34 (2006).

31. P. A. Zipfel, S. C. Bunnell, D. S. Witherow, J. J. Gu, E. M. Chislock, C. Ring, A. M. Pendergast, Role for the Abi/wave protein complex in T cell receptor-mediated proliferation and cytoskeletal remodeling. Curr Biol 16, 35–46 (2006).

32. J. C. Nolz, L. P. Nacusi, C. M. Segovis, R. B. Medeiros, J. S. Mitchell, Y. Shimizu, D. D. Billadeau, The WAVE2 complex regulates T cell receptor signaling to integrins via Abl-and CrkL-C3G-mediated activation of Rap1. J Cell Biol 182, 1231–1244 (2008).

33. T. N. Sims, T. J. Soos, H. S. Xenias, B. Dubin-Thaler, J. M. Hofman, J. C. Waite, T. O. Cameron, V. K. Thomas, R. Varma, C. H. Wiggins, M. P. Sheetz, D. R. Littman, M. L. Dustin, Opposing effects of PKCtheta and WASp on symmetry breaking and relocation of the immunological synapse. Cell 129, 773–785 (2007).

34. R. Calvez, F. Lafouresse, J. De Meester, A. Galy, S. Valitutti, L. Dupre, The Wiskott-Aldrich syndrome protein permits assembly of a focused immunological synapse enabling sustained T-cell receptor signaling. Haematologica 96, 1415–1423 (2011).

35. C. V. Carman, P. T. Sage, T. E. Sciuto, M. A. de la Fuente, R. S. Geha, H. D. Ochs, H. F. Dvorak, A. M. Dvorak, T. A. Springer, Transcellular diapedesis is initiated by invasive podosomes. Immunity 26, 784–797 (2007).

36. J. De Meester, R. Calvez, S. Valitutti, L. Dupre, The Wiskott-Aldrich syndrome protein regulates CTL cytotoxicity and is required for efficient killing of B cell lymphoma targets. J Leukoc Biol 88, 1031–1040 (2010).

37. A. T. Ritter, S. M. Kapnick, S. Murugesan, P. L. Schwartzberg, G. M. Griffiths, J. Lippincott-Schwartz, Cortical actin recovery at the immunological synapse leads to termination of lytic granule secretion in cytotoxic T lymphocytes. Proc Natl Acad Sci U S A 114, E6585–E6594 (2017).

38. B. Qu, V. Pattu, C. Junker, E. C. Schwarz, S. S. Bhat, C. Kummerow, M. Marshall, U. Matti, F. Neumann, M. Pfreundschuh, U. Becherer, H. Rieger, J. Rettig, M. Hoth, Docking of lytic granules at the immunological synapse in human CTL requires Vti1b-dependent pairing with CD3 endosomes. J Immunol 186, 6894–6904 (2011).

39. A. C. Brown, S. Oddos, I. M. Dobbie, J. M. Alakoskela, R. M. Parton, P. Eissmann, M. A. Neil, C. Dunsby, P. M. French, I. Davis, D. M. Davis, Remodelling of cortical actin where lytic granules dock at natural killer cell immune synapses revealed by super-resolution microscopy. PLoS Biol 9, e1001152 (2011).

40. P. P. Chandra, N. T. Ktistakis, in Trafficking Inside Cells: Pathways, Mechanisms and Regulation, N. Segev, Ed. (Springer, New York, 2009), chap. 11, pp. 210–232.

41. A. Chauveau, A. Le Floc’h, N. S. Bantilan, G. A. Koretzky, M. Huse, Diacylglycerol kinase alpha establishes T cell polarity by shaping diacylglycerol accumulation at the immunological synapse. Sci Signal 7, ra82 (2014).

42. X. Liu, T. M. Kapoor, J. K. Chen, M. Huse, Diacylglycerol promotes centrosome polarization in T cells via reciprocal localization of dynein and myosin II. Proc Natl Acad Sci U S A 110, 11976–11981 (2013).

43. R. Basu, Y. Chen, E. J. Quann, M. Huse, The Variable Hinge Region of Novel PKCs Determines Localization to Distinct Regions of the Immunological Synapse. PloS one 9, e95531 (2014).

44. A. Wiedemann, D. Depoil, M. Faroudi, S. Valitutti, Cytotoxic T lymphocytes kill multiple targets simultaneously via spatiotemporal uncoupling of lytic and stimulatory synapses. Proc Natl Acad Sci U S A 103, 10985–10990 (2006).

45. B. Butler, J. A. Cooper, Distinct roles for the actin nucleators Arp2/3 and hDia1 during NK-mediated cytotoxicity. Curr Biol 19, 1886–1896 (2009).

46. R. Houmadi, D. Guipouy, J. Rey-Barroso, Z. Vasconcelos, J. Cornet, M. Manghi, N. Destainville, S. Valitutti, S. Allart, L. Dupre, The Wiskott-Aldrich Syndrome Protein Contributes to the Assembly of the LFA-1 Nanocluster Belt at the Lytic Synapse. Cell reports 22, 979–991 (2018).

47. H. Ueda, M. K. Morphew, J. R. McIntosh, M. M. Davis, CD4+ T-cell synapses involve multiple distinct stages. Proc Natl Acad Sci U S A 108, 17099–17104 (2011).

48. Y. Jung, I. Riven, S. W. Feigelson, E. Kartvelishvily, K. Tohya, M. Miyasaka, R. Alon, G. Haran, Three-dimensional localization of T-cell receptors in relation to microvilli using a combination of superresolution microscopies. Proc Natl Acad Sci U S A 113, E5916–E5924 (2016).

49. J. Zhang, A. Shehabeldin, L. A. da Cruz, J. Butler, A. K. Somani, M. McGavin, I. Kozieradzki, A. O. dos Santos, A. Nagy, S. Grinstein, J. M. Penninger, K. A. Siminovitch, Antigen receptor-induced activation and cytoskeletal rearrangement are impaired in Wiskott-Aldrich syndrome protein-deficient lymphocytes. J Exp Med 190, 1329–1342 (1999).

50. L. Dupre, A. Aiuti, S. Trifari, S. Martino, P. Saracco, C. Bordignon, M. G. Roncarolo, Wiskott-Aldrich syndrome protein regulates lipid raft dynamics during immunological synapse formation. Immunity 17, 157–166 (2002).

51. E. Rivers, A. J. Thrasher, Wiskott-Aldrich syndrome protein: Emerging mechanisms in immunity. Eur J Immunol 47, 1857–1866 (2017).

52. J. H. Kim, P. Jin, R. Duan, E. H. Chen, Mechanisms of myoblast fusion during muscle development. Curr Opin Genet Dev 32, 162–170 (2015).

53. K. L. Sens, S. Zhang, P. Jin, R. Duan, G. Zhang, F. Luo, L. Parachini, E. H. Chen, An invasive podosome-like structure promotes fusion pore formation during myoblast fusion. J Cell Biol 191, 1013–1027 (2010).

54. J. H. Kim, Y. Ren, W. P. Ng, S. Li, S. Son, Y. S. Kee, S. Zhang, G. Zhang, D. A. Fletcher, D. N. Robinson, E. H. Chen, Mechanical tension drives cell membrane fusion. Developmental cell 32, 561–573 (2015).

55. M. L. Dustin, Insights into function of the immunological synapse from studies with supported planar bilayers. Curr Top Microbiol Immunol 340, 1–24 (2010).

56. L. Balagopalan, E. Sherman, V. A. Barr, L. E. Samelson, Imaging techniques for assaying lymphocyte activation in action. Nature reviews 11, 21–33 (2011).

57. E. J. Quann, E. Merino, T. Furuta, M. Huse, Localized diacylglycerol drives the polarization of the microtubule-organizing center in T cells. Nat Immunol 10, 627–635 (2009).

58. C. Fellmann, T. Hoffmann, V. Sridhar, B. Hopfgartner, M. Muhar, M. Roth, D. Y. Lai, I. A. Barbosa, J. S. Kwon, Y. Guan, N. Sinha, J. Zuber, An optimized microRNA backbone for effective single-copy RNAi. Cell reports 5, 1704–1713 (2013).

59. N. E. Sanjana, O. Shalem, F. Zhang, Improved vectors and genome-wide libraries for CRISPR screening. Nature methods 11, 783–784 (2014).

60. O. Shalem, N. E. Sanjana, E. Hartenian, X. Shi, D. A. Scott, T. Mikkelson, D. Heckl, B. L. Ebert, D. E. Root, J. G. Doench, F. Zhang, Genome-scale CRISPR-Cas9 knockout screening in human cells. Science 343, 84–87 (2014).

61. B. C. Chen, W. R. Legant, K. Wang, L. Shao, D. E. Milkie, M. W. Davidson, C. Janetopoulos, X. S. Wu, J. A. Hammer, 3rd, Z. Liu, B. P. English, Y. Mimori-Kiyosue, D. P. Romero, A. T. Ritter, J. Lippincott-Schwartz, L. Fritz-Laylin, R. D. Mullins, D. M. Mitchell, J. N. Bembenek, A. C. Reymann, R. Bohme, S. W. Grill, J. T. Wang, G. Seydoux, U. S. Tulu, D. P. Kiehart, E. Betzig, Lattice light-sheet microscopy: imaging molecules to embryos at high spatiotemporal resolution. Science 346, 1257998 (2014).

62. M. Strazza, I. Azoulay-Alfaguter, A. Pedoeem, A. Mor, Static adhesion assay for the study of integrin activation in T lymphocytes. J Vis Exp, (2014).

63. M. A. Purbhoo, D. J. Irvine, J. B. Huppa, M. M. Davis, T cell killing does not require the formation of a stable mature immunological synapse. Nat Immunol 5, 524–530 (2004).

